# Differential ADAR editing landscapes in major depressive disorder and suicide

**DOI:** 10.1101/2021.05.22.445267

**Authors:** Noel-Marie Plonski, Richard Meindl, Helen Piontkivska

## Abstract

Neuropsychiatric disorders, including depression and suicide, are becoming an increasing public health concern. Rising rates of both depression and suicide, exacerbated by the current COVID19 pandemic, have only hastened our need for objective and reliable diagnostic biomarkers. These can aide clinicians treating depressive disorders in both diagnosing and developing treatment plans. While differential gene expression analysis has highlighted the serotonin signaling cascade among other critical neurotransmitter pathways to underly the pathology of depression and suicide, the biological mechanisms remain elusive. Here we propose a novel approach to better understand molecular underpinnings of neuropsychiatric disorders by examining patterns of differential RNA editing by adenosine deaminases acting on RNA (ADARs). We take advantage of publicly available RNA-seq datasets to map ADAR editing landscapes in a global gene-centric view. We use a unique combination of Guttman scaling and random forest classification modeling to create, describe and compare ADAR editing profiles focusing on both spatial and biological sex differences. We use a subset of experimentally confirmed ADAR editing sites located in known protein coding regions, the excitome, to map ADAR editing profiles in Major Depressive Disorder (MDD) and suicide. Using Guttman scaling, we were able to describe significant changes in editing profiles across brain regions in males and females with respect to cause of death (COD) and MDD diagnosis. The spatial distribution of editing sites may provide insight into biological mechanisms under-pinning clinical symptoms associated with MDD and suicidal behavior. Additionally, we use random forest modeling including these differential profiles among other markers of global editing patterns in order to highlight potential biomarkers that offer insights into molecular changes underlying synaptic plasticity. Together, these models identify potential prognostic, diagnostic and therapeutic biomarkers for MDD diagnosis and/or suicide.

## Introduction

Depressive disorders are ranked by the World Health Organization as the largest contributor to disability and financial burden worldwide, with over 300 million people suffering depression before the start of the pandemic. In turn, depression is a major contributor to suicide deaths, with as many as 800,000 people affected yearly (Ferrari et al., 2013; World Health Organization, 2017). While treatments have been improving over the years, the disorders remain underdiagnosed, in part because our predictive power to identify future suicidality or major depressive episodes is still lacking (Kraus et al., 2019). The ongoing COVID-19 pandemic has already greatly exacerbated this burden, through psychological distress due to major pandemic disruptions of social and economic networks, as well as through direct effects of the viral infection itself, although the molecular mechanisms remain unclear (Carvalho et al., 2020; Troyer et al., 2020).

Here we propose a new approach to identifying novel neuropsychiatric biomarkers by looking at post-transcriptional modifications, more specifically, changes in RNA editing patterns by adenosine to inosine deamination catalyzed by adenosine deaminases acting on RNA (ADARs). ADAR editing is the most abundant form of RNA editing allowing for the fine-tuning of neural signaling under-pinning neural development and plasticity, and is widespread throughout the human nervous system. High-throughput studies identified over 22,000 (Z. Peng et al., 2012), and up to 147,000+ (Ramaswami et al., 2012) editing sites in the human genome, mostly located within non-coding regions. On the other hand, a small subset of experimentally confirmed editing sites (less than 1%) are located within protein coding regions and appear to have important function consequences in nerve cell ion homeostasis and signaling pathways (Hood & Emeson, 2012; Rosenthal & Seeburg, 2012). However, to the best of our knowledge, the majority of depression-related studies are focused on editing events within a single gene, mostly serotonin 2C receptor [HTR2C] (Dracheva et al., 2008; Karanović et al., 2015; Lyddon et al., 2013; Narzo et al., 2008; Schmauss, 2003, 2005; Simmons et al., 2010), with the exception of a few studies on phosphodiesterase 8A [PDE8A] (Chimienti et al., 2019; Salvetat et al., 2019) as a blood marker for neuropsychiatric disorder, and Filamin A (Stulić & Jantsch, 2013) in the frontal cortex thereby, thus, potentially missing the insights into the other players involved in synaptic plasticity.

Despite fundamental importance, one major challenge in elucidating RNA editing patterns, particularly those relevant to neural transcriptome diversity and regulation, stems from the neuron’s nuanced spatiotemporal and dynamic regulation of ADAR-editing. The majority of ADAR editing sites are found in the repetitive non-coding regions and thus may need to be filtered for studies that examine protein functional consequences. For this reason, most current studies focus on gene-centric profiles, mostly limited to a single gene or a handful of candidates at time. We propose a more global (focused on genic regions) hypothesis-generating approach, by looking at three aspects of ADAR editing (i.) across all ADAR editing sites in the entire transcriptome, (ii.) at editing events located in only the exon regions, and (iii.) at ADAR editing sites in the entire excitome. Here the excitome is defined as 92 genes containing 152 computationally predicted and experimentally validated ADAR editing sites located in ion channels, transcriptional and translational machinery and vesicle transport pathways (Supplemental Table 1) (Khermesh et al., 2016; Zhu et al., 2012). These 92 excitome genes are involved in pathways related to neuronal signaling, neurotransmitter release and recycling, which underlie synaptic plasticity (Figure 1), and include some that are implicated in neuropsychiatric disorders such as depression (Taylor et al., 2003; Ebert & Bemeier, 1996; Koenigs & Grafman, 2009), and obsessive-compulsive disorder (Zald et al., 1996).

**Figure 1:**
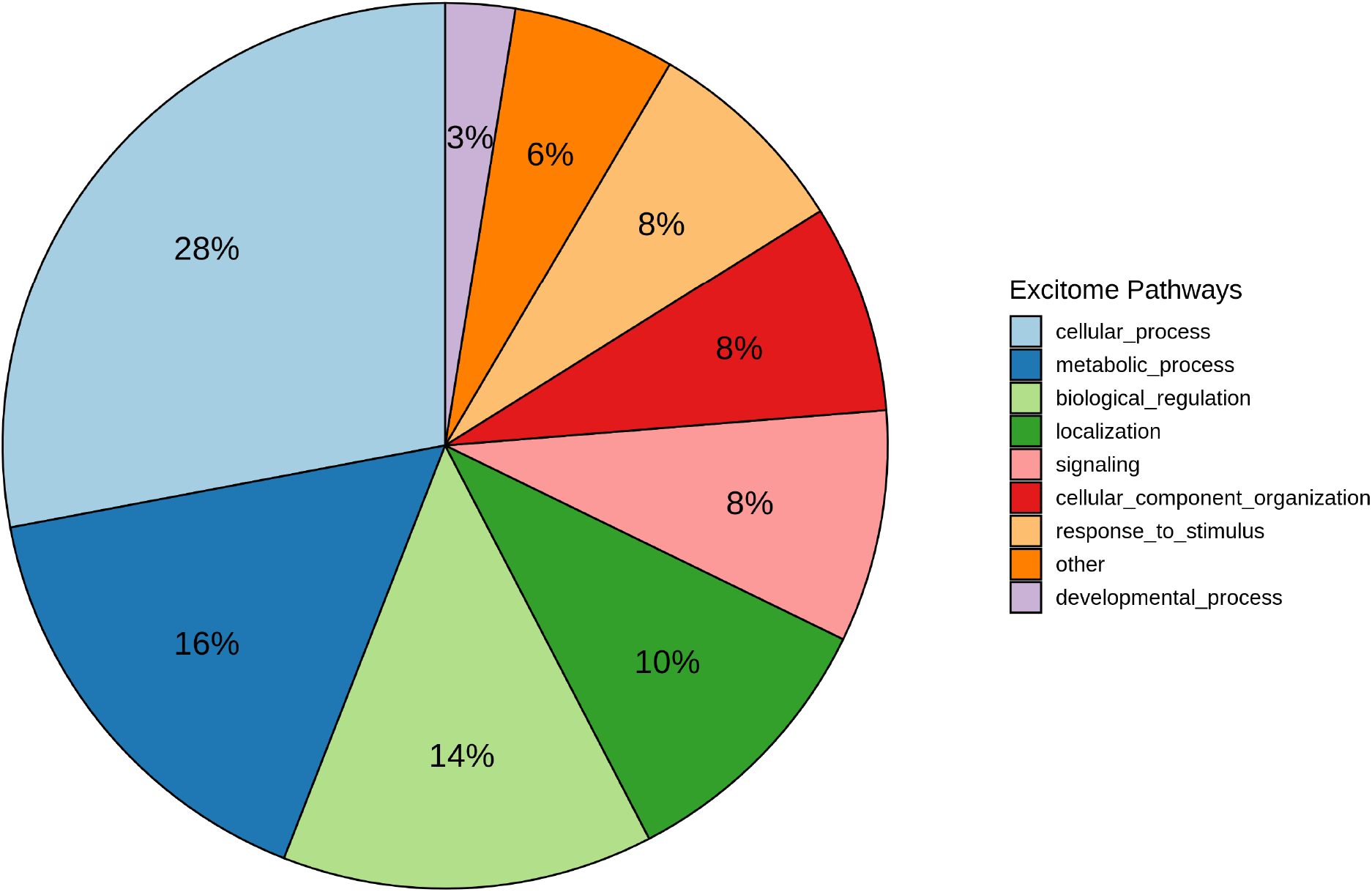
Distribution of 92 excitome genes across functional pathways.

An agnostic focus on quantifying the editing events within the genic regions and excitome as opposed to enumerating genome-wide events allows for a more comprehensive yet nuanced view of the fine-tuning of ADAR editing landscape underlying synaptic plasticity. Specifically, we examine the multidimensional complexity of the ADAR editing patterns using Guttman Scaling to reduce editing complexity to a single dimension, taking advantage of unidirectional hierarchical/chronological ordering inherent in Guttman Scale (Guttman, 1944; Proctor, 1970). Guttman scale enables us to group and rank order editing sites creating a unique profile. These chronologically ordered profiles are spatiotemporally regulated allowing for simultaneous multiple comparisons across brain regions and time points. Additionally, Guttman scale assigns a single numeric score to each sample based on these profiles, enabling targeted ranking of conditions based on the relative extent of editing across the excitome. Furthermore, assigned ranks in combination with editing frequency can then be used for machine learning-based biomarker prediction with random forest classification modeling, e.g, to identify which editing sites are associated with depression or suicidal behavior. Overall, elucidation of editing patterns at key target sites in the context of their functional pathways helps glean insight into molecular mechanisms under-pinning clinical and behavioral symptoms of neuropsychiatric disorders.

Here we use a publicly available RNA-seq data of regional postmortem brain biopsy samples to characterize patterns of transcriptome diversity, specifically, the differential ADAR editing landscapes within the excitome that are associated with fine-tuning of synaptic plasticity in neuropsychiatric disorders. We compare ADAR editing profiles between (i) suicide and natural cause of death (COD), and between (ii) diagnosis of major depressive disorder (MDD) of suicide with respect to both biological sex and brain region and their age matched controls. Our results show significant differences in ADAR editing landscapes between males and females with respect to COD and MDD, as well as between brain regions. Random forest modeling further identifies a set of potential biomarkers that could be used in conjunction with psychological assessments – to aid in suicide prevention and early intervention in MDD.

## Methods

### The RNA-seq Data

ADAR editing profiles for MDD, suicide and age-matched controls were inferred from Bioproject PRJNA394722 (Pantazatos et al., 2017), which includes post-mortem brain samples from pre-frontal cortex Brodmann’s area (BA) 9, and from Bioproject PRJNA398031 (Labonté et al., 2017), which sampled 6 different brain regions (namely anterior insula BA26 (aINS), cingulate gyrus BA25 (Cg25), dorsal lateral pre-frontal cortex (BA11 (dlPFC), nucleus accumbens (Nac), orbitofrontal cortex BA11 (OFC), and subiculum (Sub)). These studies included samples from both biological sexes, diagnosed with MDD and their age-matched healthy controls, as well as those who committed suicide with their age-matched natural death controls. There were additional 14 individuals who had an accidental COD, and thus were excluded from the final analysis due to the possibility that suicide can mistakenly be classified as accident. None of the samples had any other gross neuropathology. Furthermore, all samples tested negative for illicit psychoactive and other neurotoxic drugs and had no history of substance abuse disorders.

### Raw data analysis

Our computational pipeline, Automated Isoform Diversity Detector (AIDD) (Plonski et al., 2020) was used to map, assemble and perform variant calling on RNA-seq datasets using HISAT2 (Kim et al., 2019; Pertea et al., 2015) for alignment to GRCh37 (human reference Ensembl build release 75) annotated with both splice site and genomic single nucleotide polymorphisms (SNPs). In total, between 90-95% of reads for each of the used 303 RNA-seq samples were accurately aligned to the genome (Supplemental Table 2).

### Variant Calling

Transcriptome assembly was performed using Stringtie resulting in an aligned and annotated BAM file (Pertea et al., 2015), followed by variant calling with GATK haplotype caller (McKenna et al., 2010) to infer potential RNA editing events. The best practice settings as defined by the GATK developers as of March 2019 were used (https://software.broadinstitute.org/gatk/documentation/article.php?id=3891), except for the recommended filtering of 3 variants found within 35 bases of each other, because many of the edited sites within coding regions are clustered together (John et al., 2017) and, thus, could be filtered out as false positives. Haplotype caller is used twice in our pipeline, along with filtering steps to control for both false-positives and false-negatives. The variants already found in NCBI’s database of single nucleotide polymorphisms (dbSNP) (Sherry, 2001) were filtered out (Supplemental Table 3). However, this increases the chance that some true editing sites are being removed because some editing events have been reported as polymorphisms in dbSNP. Thus, in addition to global analysis we also used a manually curated list of excitome editing sites (Supplementary Table 1 in (Plonski et al., 2020)). SnpEff was used to predict the location and impact of variants on structure and function for global RNA editing patterns. Of these, we focus on those with high or moderate impact on protein structure and function (Cingolani et al., 2012). VCF files are available at https://www.zenodo.org/record/4140513#.YKQmUKhKguU (DOI:10.5281/zenodo.4140513).

### Statistical Analysis and Data Visualization: Global ADAR editing profiles

ADAR editing results in identification of A to G, or U(T) to C if the event occurs on the complementary strand, variants among RNA transcripts. Here we will refer to the latter category as T to C variants, keeping the genomic reference in mind. SnpEff output files are used to create matrices for nucleotide and amino acid substitutions resulting from A to G or T to C substitutions. Statistical significance of editing differences is calculated with three-way analysis of variance (ANOVA) using Tukey HSD for post-hoc testing and Cohen’s d to calculate strength of association testing. To determine the best fit model, we use Akaike information criterion (AIC) with R package “AICcmodavg” (Mazerolle 2020). This allows for us to balance variation against the number of parameters used and we select the model that explains the most variation. The models tested include full models, two-way models, three-way and all main effects all are with and without quality control measures (RNA Integrity Number (rin), potential of hydrogen (ph), postmortem interval (pmi)). Matrices of high and moderate impact editing sites are created from snpEff impact gene lists, containing the number of such sites that are then compared between conditions using differential expression analysis on the counts of sites present in each gene. PANTHER was used to perform pathway enrichment analysis using false discovery rate to calculate final p-value (Mi et al., 2021).

### Statistical Analysis and Data Visualization: Creating excitome ADAR editing landscapes

The landscapes focus on RNA editing patterns in 152 known editing sites located in the coding regions of 92 genes, listed in Supplemental Table 1 (Khermesh et al., 2016; Zhu et al., 2012), that we refer to as the excitome genes. Editing frequencies are calculated for the known editing sites in these genes by counting the nucleotides in quality-controlled aligned and assembled bam file from Stringtie-generated “stacks”. The stack depths are recorded for each excitome site. If the stack depth is greater than 10 reads, A to G (or T to C) editing frequency is determined by dividing number of Gs (or Cs) found by the total number of nucleotides found at the specific position; stacks with less than 10 reads are filtered out (Supplemental Table 4). Guttman Scaling matrices are created by assigning each site a value of 1 if editing frequency is above 0%, or 0 if editing frequency is at 0%. We also filtered out sites detected only in one of the samples as potential noise. We considered the higher cut-off of at least 5% editing frequencies inferred from each stack to reduce the chance of erroneous designation of an editing event (Sharma et al., 2019); however, the resultant lists of differentially edited genes/sites were essentially similar for each cut-off of using no cutoff (0%), 1%, or the more stringent 5%. Percent of Guttman scale rank order editing site category change between 0% and 1%, 0% and 5% and 1% to 5% cut-off values show highly similar results among editing sites (Supplemental Table 5 & Supplemental Figure 1), especially those found to be potential biomarkers (Supplemental Table 6 and Supplemental Figure 2). Thus, we will focus on results based on no cut-off threshold as to not miss editing sites with normally low editing frequency.

### Statistical Analysis and Data Visualization: Guttman Scale Analysis

Guttman Scaling is applied to rank and order the excitome ADAR editing sites based on presence of editing events while computing a score for each sample (Guttman, 1944; Proctor, 1970; Simmons et al., 2010). Custom R scripts are used with “sirt” R package to calculate probabilistic Guttman Scale model. The samples are scored based on the hierarchal ranks and a three-way ANOVA included in R-base is used to determine statistical differences based on scores between the cause of death and MDD diagnosis. The rank order of editing sites is used to create topological editing profiles that can then be compared between conditions to see variation in editing landscapes. Rank groups can be additionally grouped into three categories within each condition. Specifically, the first rank group category consists of sites that are completely edited in all samples. Category two includes sites that have both edited and unedited samples. The last rank group category contains no editing in any sample. These assigned Guttman categories are compared within brain region and sex between COD and MDD diagnosis using the “UpSetR” package to create upset plots to explore editing site variation among regions and sexes (Conway et al., 2017; Lex et al., 2014).

### Statistical Analysis and Data Visualization: Random Forest classification modeling

To identify potential biomarkers of MDD or suicide, we applied random forest classification modeling using “randomForest” R package which implements Breiman’s random forest algorithm (Liaw & Wiener, 2002) enabling us to integrate the editing markers with other transcriptomics and clinically relevant indicators (Supplemental Figure 3). The measurements found to have significant differences – including the Guttman scale maximum likelihood estimates of scores, excitome editing frequencies, snpEff predicted with high and moderate impact on protein structure or function, counts of editing in exon regions, previous MDD diagnosis, biological sex, and brain regions – were used as variable parameters in random forest modeling. The novel approach of quantifying Guttman scaling and then using a non-parametric alternative, which utilizes bootstrap consensus of random classification decision trees (random forest), enables us to predict clinically relevant biomarkers using Variance Importance (VIMP) and Minimal Depth (Ishwaran & Kogalur, 2010) cut-offs. For more on parameter determination for modeling see supplemental methods.

## Results

The original analyses conducted on the RNA-seq data sets (Pantazatos et al., 2017; Labonté et al., 2017) focused on differential gene and exon expression, rather than ADAR editing, in MDD and suicide. Both studies identified differential expression of genes involved in neural pathways (such as oligodendrocyte differentiation, regulation of glutamatergic neurotransmissions and oxytocin receptor expression, DNA-dependent ATPase) and immune-related genes linked with altered microglia and/or astrocyte activity, resulting in changes to synaptic plasticity and neuroprotection. Although variant calling was performed in one study (Pantazatos et al., 2017), its focus was on identifying genomic single nucleotide polymorphisms and indels as opposed to our approach of investigating RNA editing landscapes. The second study (Labonté et al., 2017) compared differential gene expression between biological sexes across brain regions and did not examine variants. Based on their findings of significant sex differences, here we treat male and female samples separately to compare differential editing patterns between sexes.

We compared global ADAR editing landscapes across COD, and MDD diagnosis, by gender and brain region. There were no significant changes to global ADAR editing profiles between COD, brain regions, or MDD, but there were statistically significance differences when compared by gender with a small effect size (Cohen’s d = 0.202846813). However, there were relatively small sample sizes across categories, with some having rather large variances. Quality control measures ph was also significant with rin and pmi approaching significance, therefore they were included in the model (Table 1). When further exploring the differential trends in global editing patterns between sexes, there are trends in global editing patterns such as females show more ADAR editing with suicide as the COD in the dlPFC including BA9, aINS, and Cg25 (Figure 1A), whereas males show decreased editing in these regions (Figure 1B). Additionally, females showed decreased editing in OFC, NAcc, and sub and males also had decreased editing in the sub but had increased editing in OFC, no change in editing in the NAcc. Females also have increased editing in the OFC with MDD diagnosis but not with suicide as the COD (Supplemental Figure 4A). The majority of these editing sites, over 75%, are located in introns in all brain regions, males and females with no significant differences between COD or MDD diagnosis (Supplemental Table 7 & Supplemental Figure 5). Up and down stream regions have over 10% and 8% respectively with less than 2% located in exons. MDD diagnosis and COD in males shows increased editing in the OFC but PFCBA9 along with the other brain regions shows decreased editing (Supplemental Figure 4B). Females like males have decreased editing in PFCBA9, the NAcc, and aINS unlike with females who commit suicide who have increased in editing in PFCBA9 and aINS. Females have decreased editing in Sub in those that commit suicide, while males showed increased editing. Females have increased editing in the PFC and Cg25 with MDD diagnosis while males have decreased editing in these regions. Changes to editing patterns in these specific regions can potentially explain the broad spectrum of symptoms seen in depression and suicidal behavior. Moreover, the differences between editing profiles in biological sexes can contribute to differences in manifested symptoms. While these findings offer an initial glimpse into potentially distinct editing patterns associated with different diagnoses, further extensive validation is needed to link specific editing events to clinical and/or behavioral symptoms.

**Table 1:**
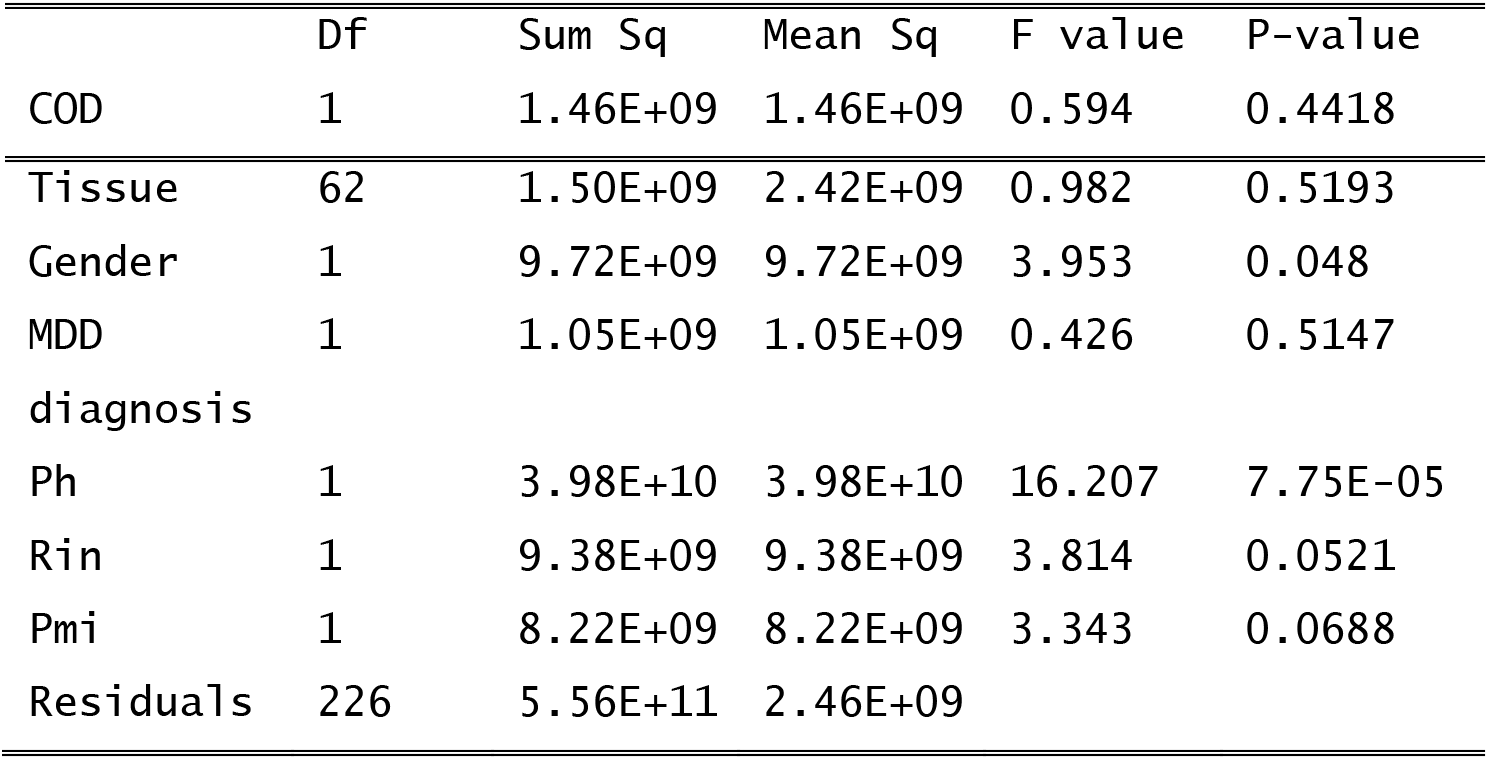
Four-way ANOVA with all four main effects and co-variant quality control measures ph, rin, pmi for ADAR global editing.

|  | Df | Sum Sq | Mean Sq | F value | P-value |
| --- | --- | --- | --- | --- | --- |
| COD | 1 | 1.46E+09 | 1.46E+09 | 0.594 | 0.4418 |
| Tissue | 62 | 1.50E+09 | 2.42E+09 | 0.982 | 0.5193 |
| Gender | 1 | 9.72E+09 | 9.72E+09 | 3.953 | 0.048 |
| MDD | 1 | 1.05E+09 | 1.05E+09 | 0.426 | 0.5147 |
| diagnosis |  |  |  |  |  |
| Ph | 1 | 3.98E+10 | 3.98E+10 | 16.207 | 7.75E-05 |
| Rin | 1 | 9.38E+09 | 9.38E+09 | 3.814 | 0.0521 |
| Pmi | 1 | 8.22E+09 | 8.22E+09 | 3.343 | 0.0688 |
| Residuals | 226 | 5.56E+11 | 2.46E+09 |  |  |

Ward’s test using the number of predicted high or moderate impact editing sites in individual genes was done using DESeq2 differential expression approach, identifying 660 genes with increased number of ADAR editing sites (Supplemental Table 8). These moderate/high impact editing sites included non-synonymous amino acid changes, and editing in 5’ or 3’ prime untranslated regions with anticipated effects on translational activity. The top 5 genes included SSH2 (slingshot protein phosphatase 2) involved in axonal guidance, TRIM44 (tripartite motif containing 44) important in differentiation and maturation of neural cells, GMR7 (glutamate metaboltrophic receptor 7) involved in excitatory signaling, NRCAM (neuronal cell adhesion molecule) responsible for axonal cone growth, and TJP1 (tight junction protein 1) (Figure 3A). These and other differentially edited genes were identified as part of the following gene ontology pathways: cell junction assembly, organelle assembly, mitotic cell cycle, metabolic process (Figure 3B). Pathway enrichment showed three-fold increase in genes involved in cell junction assembly, a two-fold increase in genes involved in organelle assembly, and a one-fold increase in those involved in miotic cell cycle, and metabolic process (Figure 3C).

**Figure 2:**
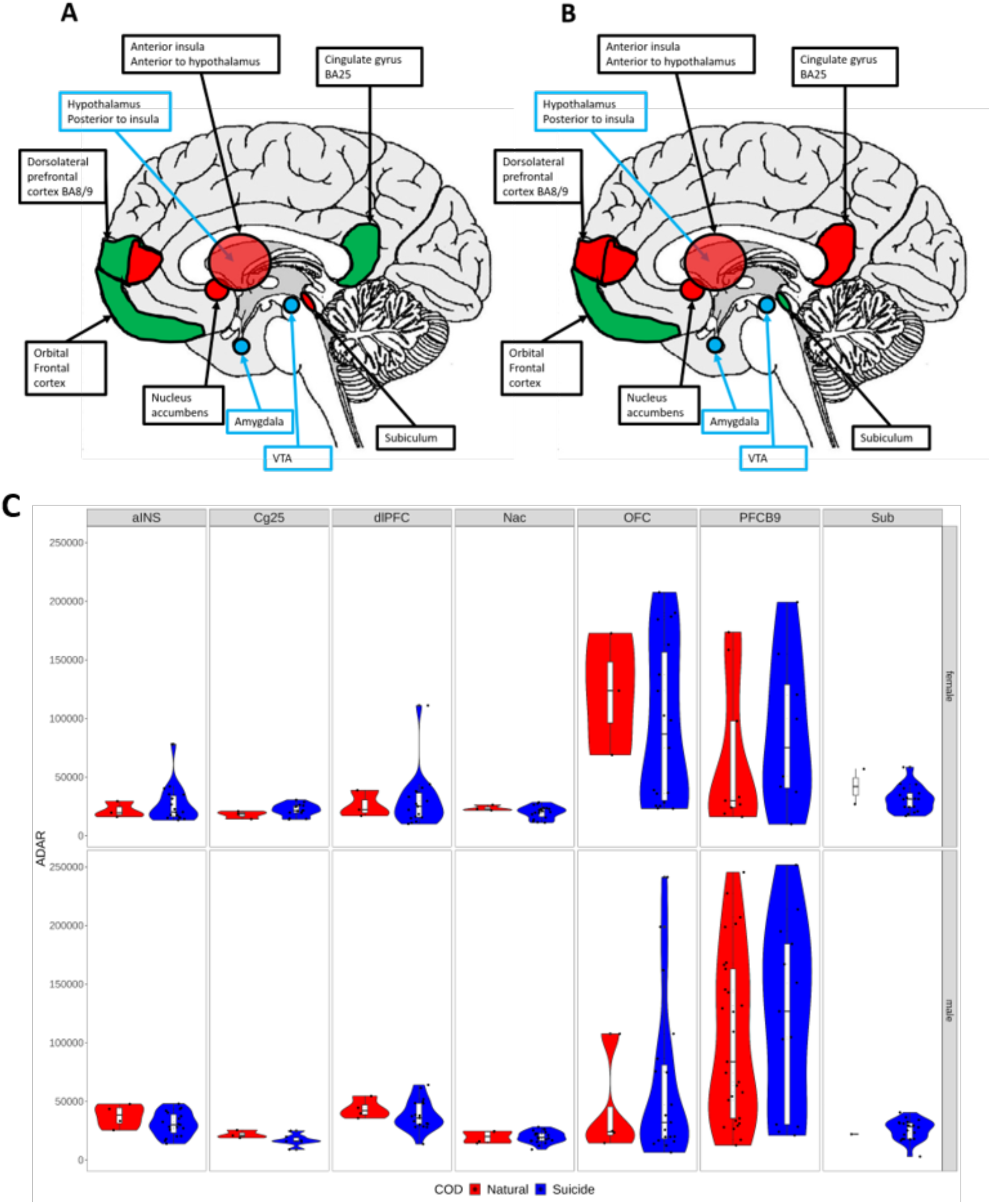
Patterns of global ADAR editing changes across natural and suicide causes of death (COD) in females (A) and males (B). Increases in editing are shown in green, decreases in editing are shown in red, and no change is shown in yellow. Reference anatomy features shown in blue helps in the location of brain regions and are involved in pathways within studied regions. (C) Global differential ADAR editing patterns in suicide vary between brain regions in females and males, as well as between suicide and natural causes of death (shown in blue and red in violin plots, respectively).

**Figure 3:**
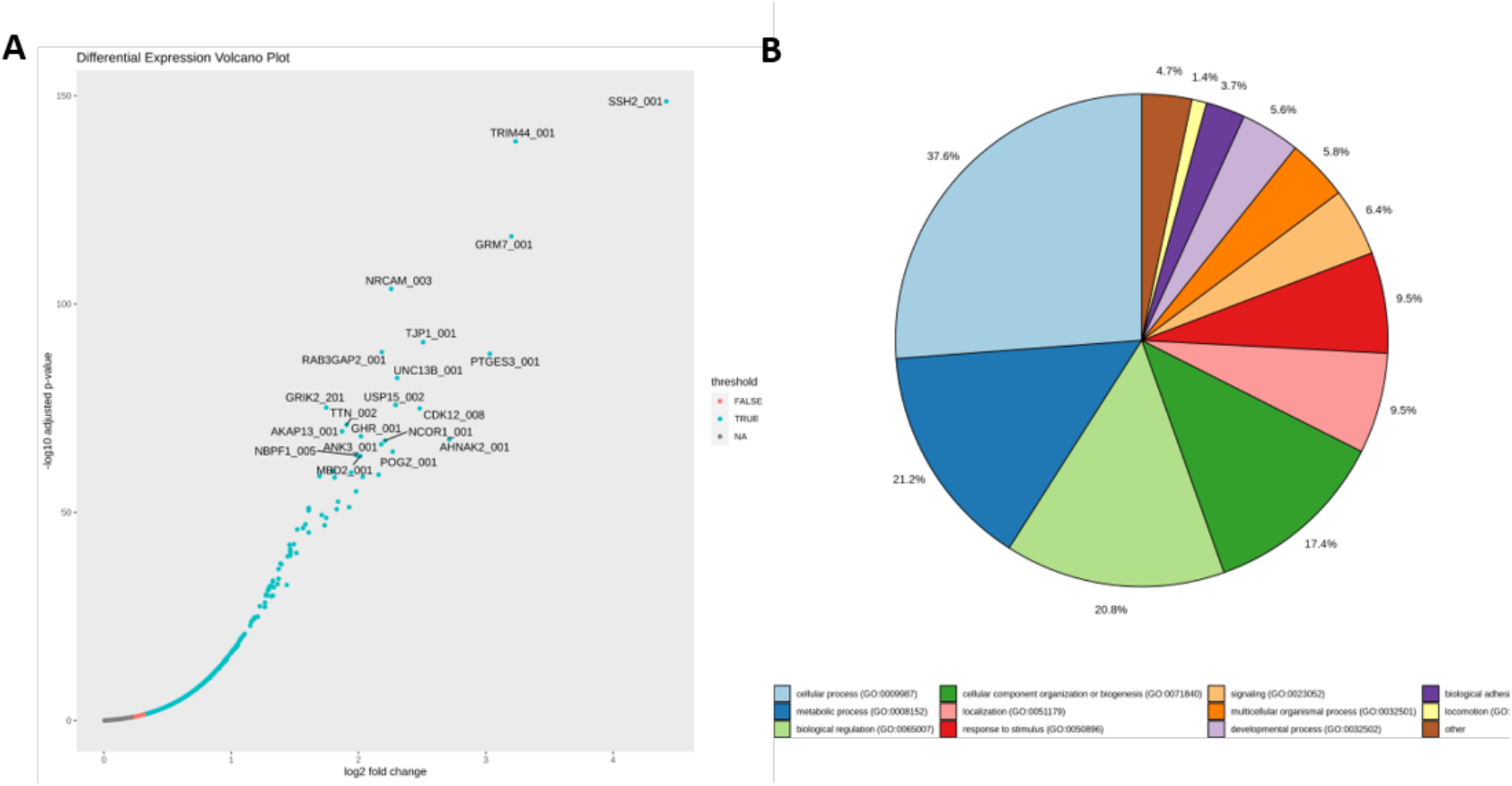
Distribution of differentially edited moderate impact editing sites pathways. 660 genes with increased editing are shown in the volcano plot (A), with significantly edited genes shown in blue and non-significant ones in red, per with FDR p-value of 0.05. The percentages of genes found in 9 gene ontology pathways (B).

Changes to these pathways can affect the neural cells communication with other cells by changing ability to package neurotransmitters and form junctions with other nearby cells, thereby leading to changes in synaptic plasticity.

### Mapping ADAR editing landscapes with Guttman Scale

Because of its relevance to synaptic plasticity, the excitome editing needs to be considered as a separate subset of global editing sites for us to better understand plasticity changes seen in depressive disorders. Guttman scaling maximum likelihood scores using the excitome editing sites were significantly different in COD and tissue with MDD diagnosis approaching significance and no significant difference between genders (Table 2). COD had a strong effect size and tissue had 7 statistically significant tissue pairs among the 21 pairs tested, with two of the comparisons having moderate effect size and four of the comparisons having high effect size showing significantly large differences with respect to Guttman Scale scoring (Table 3). Looking at the excitome compared to global ADAR editing patterns females had higher scores indicating increased ADAR editing in all brain regions in suicide except in NAcc (Figure 4A). Males had increased editing in PFC including BA9, NAcc, and sub and decreased scores in OFC, aINS, and Cg25 in suicide (Figure 4B). Females diagnosed with MDD had similar patterns to females who committed suicide except in the PFC and sub where there are lower scores or less editing, and increased scores in the NAcc (Supplemental Figure 6A). Males diagnosed with MDD have similar increases as seen in females diagnosed with MDD but different patterns than those seen in males with suicide (Supplemental Figure 6B). There were increases in scores in all regions except PFCBA9 in males and females diagnosed with MDD. Additionally, females show decreases in editing in the PFCBA8.

**Figure 4:**
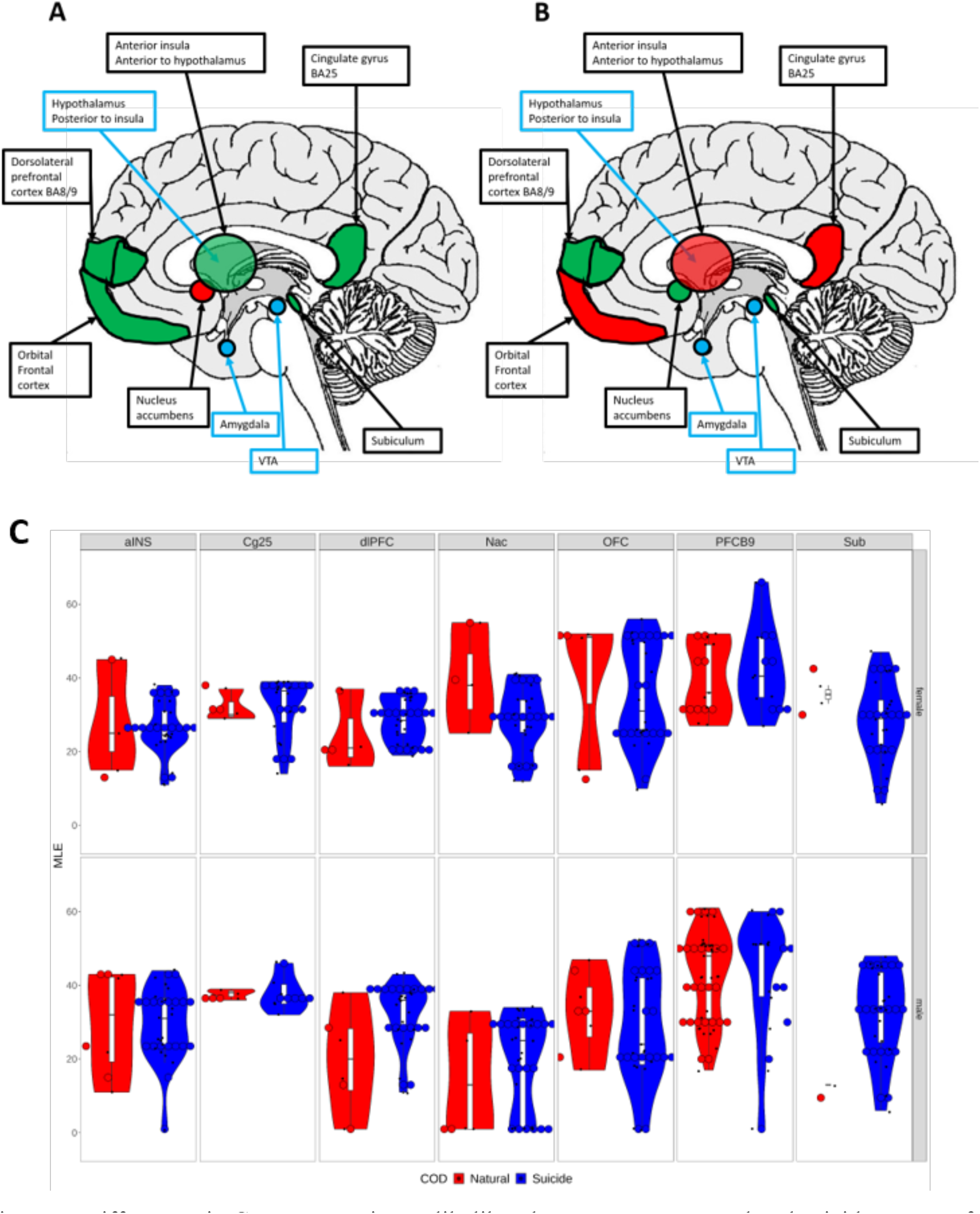
Differences in Guttman maximum likelihood scores across natural and suicide causes of death (COD) in females (A) and males (B). Differential ADAR editing sites found in Guttman scale MLE scores in suicide are highlighted, with increases in editing shown in green, decreases in editing shown in red, and no change shown in yellow. Reference anatomy features shown in blue helps in the location of brain regions and are involved in pathways within studied regions. Similar editing landscapes are observed between females and males who commit suicide, with differences seen in Cg25 and OFC regions. Males who commit suicide show a different ADAR editing pattern than those of any of the other categories. Similarly to the global differential ADAR editing patterns, Guttman scale MLE scores vary between brain regions in females and males in suicide, as well as between suicide and natural causes of death (shown in blue and red in violin plots, respectively). Violin plot shows spread of global ADAR editing sites among samples with red representing suicide and blue natural cause of death (C).

**Table 2:** Four-way ANOVA with all four main effects and co-variant quality control measures ph, rin, pmi for Guttman Scaling scores.

|  | Df | Sum Sq | Mean Sq | F value | P-value |
| --- | --- | --- | --- | --- | --- |
| COD | 1 | 1511 | 1510.6 | 11.396 | 0.000866 |
| Tissue | 6 | 18445 | 297.5 | 2.244 | 8.55E-06 |
| Gender | 1 | 20 | 19.9 | 0.15 | 0.69896 |
| MDD | 1 | 362 | 361.8 | 2.729 | 0.0999 |
| diagnosis |  |  |  |  |  |
| Ph | 1 | 59 | 59.5 | 0.449 | 0.503725 |
| Rin | 1 | 0 | 0 | 0 | 0.985178 |
| Pmi | 1 | 160 | 160.1 | 1.208 | 0.272883 |
| Residuals | 226 | 29958 | 132.6 |  |  |

**Table 3:** Significant differences and strength of association for Guttman Scaling scores.

|  | Difference Between<br>Mean | Lower<br>CI | Upper<br>CI | P adj | Cohen's d |
| --- | --- | --- | --- | --- | --- |
| Suicide/Natural | -5.101597 | -8.210373 | -1.992785 | 0.0013803 | 0.44303 |
| PFCB9/Nac | 15.25231 | 8.322594 | 22.18203 | 0 | 1.578317 |
| PFCB9/aINS | 11.43932 | 4.421666 | 18.45698 | 4.31E-05 | 1.246225 |
| PFCB9/dlPFC | 10.1666 | 3.148938 | 17.18425 | 0.000459 | 1.135699 |
| Sub/PFCB9 | -8.74082 | -15.9567 | -1.52497 | 0.006876 | 1.057739 |
| OFC/Nac | 8.608444 | 1.179246 | 16.03764 | 0.011725 | 0.748538 |
| Nac/Cg25 | -9.84967 | -18.4939 | -1.20543 | 0.014167 | 0.88381 |

To further explore differences in editing landscapes, we looked at the number of changes to editing patterns reflected by Guttman hierarchical rank order, where the smaller magnitude of change indicates higher homogeneity/similarity across phenotypes. Regardless of brain region or sex, 7 editing sites did not change their rank positions, indicating that these sites likely do not play a role in COD. There were 22 sites that decreased no more than 5 ranks from natural cause of death to suicide, and 15 sites that increased no more than 5 ranks, respectively (Figure 5A), likely indicating that these are relatively minor players, if any, in suicide. On the other hand, two-thirds of the editing sites were more than 5 ranks away from where they were positioned in natural COD, signifying major heterogeneity across samples with the same phenotype, making them good candidates for follow-up differential diagnostic studies. Similarly, there were 25 sites that decreased no more than 5 ranks from control to MDD diagnosis and 39 sites that increased no more than 5 ranks (Supplemental Figure 7A), unlikely to be major players. On the other hand, three-fourths of the editing sties were more than 5 ranks away from control samples compared to those with MDD diagnosis, indicating these editing sites may contribute to molecular mechanisms behind symptoms of depression. Overall, when differences between brain regions and sexes within suicide and MDD diagnosis are considered, both disordered phenotypes have a broader and higher variation in ranking of editing sites than that of those with natural causes of death, or age matched controls without a MDD diagnosis (Figure 5B & Supplemental Figure 7B). This observed editing diversity may be contributing to the broad spectrum of manifested symptoms seen in depression and suicidal behavior.

**Figure 5:**
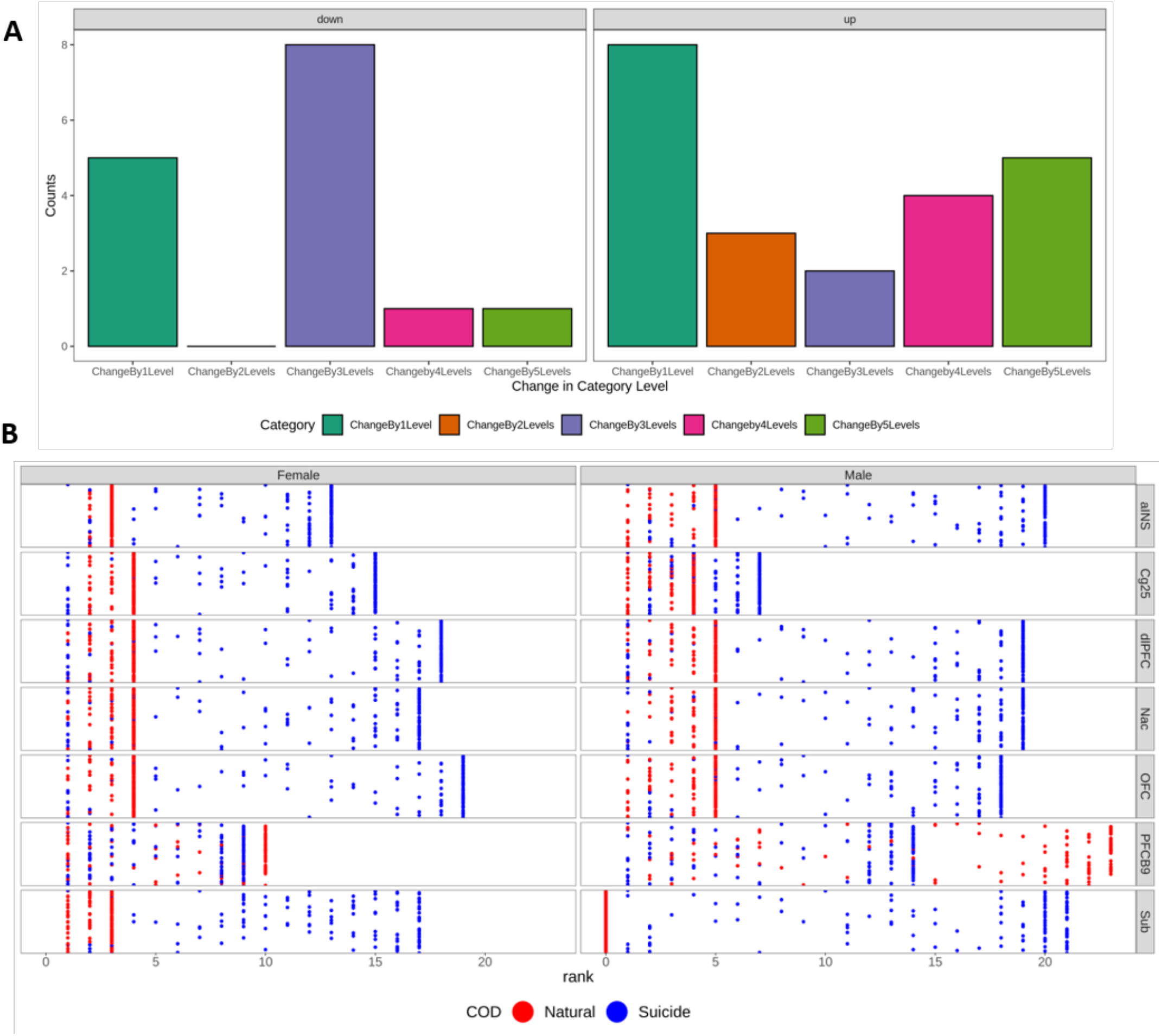
Changes in Guttman scaling rank order determined by editing status of each site in suicide and natural COD. Number of genes that change in Guttman rank order, whether up or down, from one rank change to five rank changes (A). Guttman Scaling assigned rank orders show more variation in suicide (shown in blue) than natural cause of death (shown in red) in all brain regions and in both genders indicating more editing in the excitome in suicide then in natural cause of death (B)

To further explore changes to hierarchical rank order we assigned categories based on editing status allowing us to focus on overall changes seen in suicide between brain regions and sex. The Sub region in females has almost double the amount of editing changes than any other region, 80 sites, with 11 of those being unique to the region (Figure 6). PFCB9 in females and NAcc in males have 2 unique editing sites changing in the region. The dlPFC, NAcc and OFC in females have only 1 unique to each region. dlPFC in females has one less editing site than in males however females have one unique editing site change and males have no unique changes in the region. aINS and Cg25in both males and females and dlPFC and OFC in males have no unique editing sites to just that region. The Sub of males in MDD diagnosis had the most changes to editing sites, 49 sites, with the most unique editing sites, 8 sites. PFCB9 in females and Cg25 in males has 3 unique editing site changes. NAcc and PFCB9 in males had two unique editing sites. The rest of the regions had 1 unique editing site to each region except the OFC and dlPFC in males which have no unique changes in editing sites (Supplemental Figure 8). The low number of intersections in most regions indicates high variation in the specific editing sites across brain regions. This suggests that any attempts to link changes at editing sites with specific behavioral symptoms in future experiments should potentially focus on subsets of different editing targets depending on the diagnosis and/or brain region of interest.

**Figure 6:**
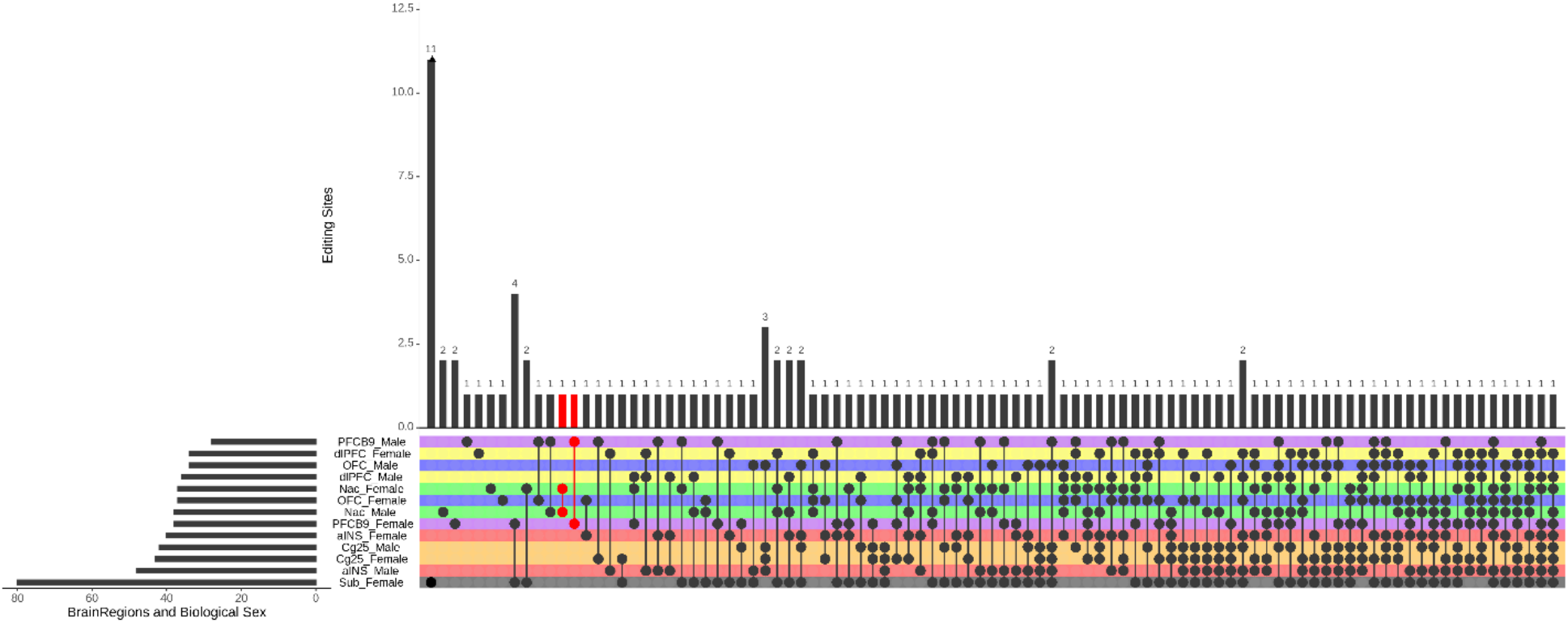
Upset plot of genes within regions and sexes that shared similar changes in Guttman rank category editing status in COD. The upset plot (Lex et al., 2014) shows each region in a different color, with each sex showing matching colors. The bars to the right show how many editing sites have changes to category, in order from the least amount of editing changes on the top with the most on the bottom. The bars across the top show how many editing sites are in each intersection. The dots in the colored bars represent intersections of the same editing sites in the region the dots are connected too. The connected dots shown in red represent the same region where both sexes share the same editing sites. For example, the first dots in red represent 1 editing in common between females and males in Nac which has 40 changes to editing sites in both males and females. The second red dots represent 1 editing site in common between females and males in the PFCB9. Larger number of dots under the bar indicate higher number of regions that share changes in that editing site.

### ADAR editing biomarkers prediction with random forests to predict biomarkers

Random forest classification modeling resulted in strong prediction of COD and moderate prediction of MDD diagnosis. The COD training model had an accuracy of 85% percent with area under the curve (AUC) of 91.7, and the COD test set model had 91% accuracy with AUC 97.9 (Table 4). The MDD training model had an accuracy of 72% with AUC of 79.7, and the test set model was 75% accurate at predicting MDD diagnosis with AUC 82.5 (Table 4). These models were then used to predict potential clinically relevant biomarkers for diagnosis, prognosis, appropriate treatments and potential targets for drug therapy development by finding the variables most important to the COD or MDD diagnosis prediction

**Table 4:** Random Forest Model validation of both training and test dataset for MDD diagnosis and COD. The average number of nodes needed to create trees with other measures of accuracy of used to use the tree for prediction. These models are both accurate and precise with low overall error rates for prediction of biomarkers.

|  | TrainingMDD | TestMDD | TrainingCOD | TestCOD |
| --- | --- | --- | --- | --- |
| Sample size | 242 | 40 | 242 | 48 |
| Average no. |  |  |  |  |
| nodes | 16.68 | 16.68 | 9.292 | 9.292 |
| Accuracy | 0.72314 | 0.75 | 0.855372 | 0.916667 |
| misclassification | 0.27686 | 0.25 | 0.144628 | 0.083333 |
| true pos rate | 0.876812 | 0.8 | 0.917582 | 0.941176 |
| false pos rate | 0.480769 | 0.3 | 0.333333 | 0.142857 |
| true neg rate | 0.519231 | 0.7 | 0.666667 | 0.857143 |
| precision | 0.707602 | 0.727273 | 0.893048 | 0.941176 |
| prevalence | 0.570248 | 0.5 | 0.752066 | 0.708333 |
| null error rate | 0.429752 | 0.5 | 0.247934 | 0.291667 |
| AUC | 79.73 | 82.5 | 91.7 | 97.9 |
| Test set error |  |  |  |  |
| rate | 0.28,0.48,0.12 | 0.25,0.3,0.2 | 0.14,0.33,0.08 | 0.08,0.14,0.06 |

Random forest modeling predicted top scoring biomarkers found in common between both VIMP/minimal depth (Supplemental Figure 9) and partial dependence analysis (Supplemental Figure 10) can be used to provide insight into the molecular mechanisms underlying suicidal behavior (Table 6). Further, we examined differential editing frequencies of the top biomarkers. Of these top biomarkers, IGFBP7(position 95, here and later) had no significant difference between COD, MDD diagnosis, gender or brain region but there was a significant difference for COD when brain region were also considered looked at with a strong strength of correlation in 11 out of 21 comparisons between tissues (F=5.411, p=0.000104, Table 6). NOVA1(387) showed no significant difference between COD, MDD diagnosis, or brain region but approached significance between genders (F=2.749, p=0.0987). PCP4(42) was signification different in COD (F=8.797, p=0.003338, Cohen’s d = 0.149405) and brain region (F=1.966, p=0.000192) but not MDD diagnosis or gender. VWF(789) is statistically significant with extremely high strength of association between tissues (F=1.940, p=0.000237, Table 6) but not COD, gender or MDD diagnosis. VWF(484) also was statistically significant between tissues with 5 out of 21 comparisons have strong strength of correlation (F=1.425,p=0.0338, Table 6) but not between COD, MDD diagnosis, or gender. Another editing site VWF(789), was significantly different for tissues with a moderate strength of association for 5 out of the 21 tissue comparisons, but not for COD, MDD diagnosis or biological sex (Table 6). This further supports the identification of some of these as potential biomarkers for both suicide risk and depression. Overall, editing site frequencies in males had less editing in suicide compared to controls, whereas females have more editing in most brain regions. In addition to editing site frequencies, variables such as transcript-level high impact on protein structure and function, brain region, maximum likelihood scores from Guttman scale, and global ADAR editing counts were also in the top 30 variables for both the COD and MDD random forest models. Notably, variables such as the number of ADAR editing sites found in the exon region of the transcript, gene-level high impact counts and transcript-level moderate impact counts were part of the top 30 for the MDD, but not COD diagnosis model, indicating editing sites in the protein coding regions are more likely to be underlying behavioral symptoms seen in depressive disorders but not the pathology of suicide.

**Table 5:** List of potential biomarkers as determined by our statistical model. ADAR editing sites are identified by the amino acid changes at respective codon positions.

| Gene Name | GeneID | ADAR editing site |
| --- | --- | --- |
| signal recognition protein 9 | SRP9 | S75G |
| insulin growth factor binding protein 7 | IGFBP7 | K95R |
| filamin A | FLNA | Q2341R |
| von Willebrand factor | VWF | H484R, Q852R, T789A, T1381A |
| Nei Like DNA Glycosylase 1 | NEIL1 | K242R |
| NOVA Alternative Splicing Regulator 1 | NOVA1 | S387G |
| acid phosphatase 1 | ACP1 | Q106R |
|  | PCP4 | A42A |
| Gamma-aminobutyric acid type A receptor subunit<br>alpha 3 | GABRA3 | I342M |
| Glutamate ionotropic receptor kainate type subunit 2 | GRIK2 | I567V |

**Table 6:** Significant differences between tissue pairs and strength of association for editing site frequencies.

| ADAR<br>editing sites | Tissue<br>Comparisons | Difference<br>in means | Lower<br>CI | Upper CI | p adj | Cohen's<br>d |
| --- | --- | --- | --- | --- | --- | --- |
| IGFBP7_95 | PFCB9-aINS | 25.6017 | 17.5762 | 33.62719 | 0 | 2.900758 |
| IGFBP7_95 | PFCB9-dlPFC | 25.39508 | 17.36959 | 33.42058 | 0 | 2.882774 |
| IGFBP7_95 | PFCB9-Nac | 25.61167 | 17.68674 | 33.53661 | 0 | 2.900758 |
| IGFBP7_95 | PFCB9-OFC | 19.12163 | 11.09613 | 27.14713 | 0 | 2.33674 |
| IGFBP7_95 | Sub-Nac | 14.47529 | 5.764733 | 23.18585 | 2.78E-05 | 1.599022 |
| IGFBP7_95 | Sub-aINS | 14.46531 | 5.663158 | 23.26747 | 3.59E-05 | 1.139177 |
| IGFBP7_95 | Nac-Cg25 | -16.1292 | -26.0149 | -6.24355 | 4.22E-05 | 1.452813 |
| IGFBP7_95 | Sub-dlPFC | 14.2587 | 5.456547 | 23.06086 | 4.95E-05 | 1.121194 |
| IGFBP7_95 | Cg25-aINS | 16.11925 | 6.152772 | 26.08573 | 5.13E-05 | 1.566699 |
| IGFBP7_95 | dlPFC-Cg25 | -15.9126 | -25.8791 | -5.94616 | 6.79E-05 | 1.43483 |
| IGFBP7_95 | Sub-PFCB9 | -11.1364 | -19.3885 | -2.88423 | 0.001507 | 1.761581 |
| VWF_484 | PFCB9-Nac | 1.31E+01 | 3.167921 | 22.97107 | 0.002114 | 2.900758 |
| VWF_484 | PFCB9-aINS | 1.31E+01 | 3.041365 | 23.09581 | 0.002549 | 2.900758 |
| VWF_484 | PFCB9-dlPFC | 1.31E+01 | 3.041365 | 23.09581 | 0.002549 | 2.882774 |
| VWF_484 | PFCB9-OFC | 1.31E+01 | 3.041365 | 23.09581 | 0.002549 | 2.33674 |
| VWF_484 | PFCB9-Cg25 | 1.38E+01 | 1.92354 | 25.62315 | 0.011314 | 1.447944 |
| VWF_789 | PFCB9-dlPFC | 10.98844 | 2.519996 | 19.45689 | 0.002717 | 2.882774 |
| VWF_789 | PFCB9-OFC | 10.79905 | 2.330603 | 19.26749 | 0.003479 | 2.33674 |
| VWF_789 | PFCB9-aINS | 10.38238 | 1.913935 | 18.85083 | 0.005898 | 2.900758 |
| VWF_789 | Sub-PFCB9 | -10.0596 | -18.7672 | -1.35202 | 0.012133 | 1.761581 |
| VWF_789 | PFCB9-Cg25 | 11.32875 | 1.32105 | 21.33645 | 0.015227 | 1.447944 |

## Discussion

Studies have found differences in gene expression during different stages of MDD as well as remission when on drug therapy (Hepgul et al., 2013; Mamdani et al., 2011). In addition, studies have also found differential gene expression in suicide attempts (Flory et al., 2017) and in completed suicides (Cabrera et al., 2019). Our findings of differential editing in these regions offer novel insights into patterns of differential gene regulation and function that can be driving synaptic plasticity across multiple pathways in the brain. Differential editing landscapes in any of the regions studied here can potentially explain molecular mechanisms behind diverse clinical behaviors and symptoms. In addition to focusing on global editing pattern, our unique approach of using Guttman scaling combined with random forest modeling allows for us to highlight biomarkers from the top scoring editing sites. These editing biomarkers give insight into potential mechanistic links to the fine-tuning of synaptic plasticity and neuropsychiatric disorders.

In particular, our results identified significant differences in editing profiles across different brain regions, where each brain region examined (aINS, OFC, dlPFC, Cg25, aINS, NAcc, Sub, dlPFC9) exhibited distinct editing profiles between controls and COD or MDD diagnosis. These regions have been linked to a variety of neurological and psychiatric disorders, symptoms and manifestations. Editing changes in target genes in the aINS is part of many pathways in the brain and contribute to the molecular underpinnings of symptoms of depression and suicidal behavior including changes to emotional homeostasis and risk assessment (Beauchaine et al., 2019; Soloff et al., 2012; Gogolla, 2017). Differential editing profiles in the OFC can alter neurotransmitters in emotion and reward pathways (Kolb et al., 2003). Structural abnormalities (W. Peng et al., 2019), and connectivity disturbances (Rolls et al., 2020) have been noted in the OFC depression and anxiety. Editing events changing protein structure or function can contribute the morphological and molecular dysfunctions seen in this region. Working memory, cognitive flexibility and planning are part of the function of the PFC (Duncan, 2001; Miller & Cohen, 2001) and disturbances here are implicated in depression (Drevets et al., 1998), therefore differential editing profiles can help explain on a molecular level changes resulting in suicidal behaviors. Changes in blood flow and metabolic function in the Cg25 been implicated in depression (Ebert & Bemeier, 1996), and we have identified editing targets present in depression and suicide that are linked with those functions. Behaviorally, differential edits at these regions can result in changes in mood and self-esteem (Vogt, 2005). Deep brain stimulation in the Nac can help with treatment resistant depression (Bewernick et al., 2010), as interneurons neurons located here help regulate depression (Warner-Schmidt et al., 2012). Calcium channels like the ones we identified as differentially edited in depression and suicide may control depressive like behaviors (Goffer et al., 2013).

Cell junction assembly and organization were two pathways that were overrepresented over two-fold. Cell junction formation and control is important for neuronal cell synaptic formation behind neural plasticity. Increases to ADAR editing in the genes involved in this pathway can alter the neuron’s ability to form new synaptic functions or remodel existing synapses causing changes to synaptic plasticity in suicidal behavior (Goda & Davis, 2003; Li & Sheng, 2003). Microtubule process and organization is important for neural development (Lasser et al., 2018), and neuron structure and function in the dopaminergic function especially in depression (Yang et al., 2018). Cell component assembly, biogenesis, and organization and organelle formation all play a role in neurotransmitter transport and release, and we have shown significant increases in the amount of editing found in genes in these pathways in both suicide and depression. Organelles are needed for proper packaging, trafficking, holding and tracking of neurotransmitters for cell signaling that drives post synaptic plasticity (Kennedy & Ehlers, 2006). Metabolic function pathways are also overrepresented with higher frequency of editing in genes in these pathways. Metabolic function is important in not only normal housekeeping functions of cells, but also providing energy for cellular processes involved in signaling and synaptic plasticity. The importance and variety of these pathways may be behind the wide range of behavior and somatic symptoms seen in suicidal behavior.

Furthermore, the top potential biomarkers predicted in our study offer insight into the specific molecular mechanisms under-pinning MDD and suicidal behavior. Changes to editing in SRP9 affect protein localization (Ivanova et al., 2015), which in turn can affect not only the cells’ ability to function at a normal level, but also package and deliver neurotransmitters for synaptic plasticity. Furthermore, differential editing can affect proteolytic degradation, angiogenesis and induce apoptosis by altering spatiotemporally regulated variant isoform of IGBPF7 another potential biomarker (Godfried Sie et al., 2012). Editing in FLNA which interferes with many proteins can change cell motility and signaling ability by affecting the neurons’ ability to form the proper microskeleton used for stability, transport, nuclear division and forming connections to other cells this can result in the ability to control blood pressure (Jain et al., 2018) and for proper neurodevelopment (Eran et al., 2013). The two differential editing sites we found in VWF can affect the link between the endothelium of blood vessels and platelets (Randi et al., 2018). Stress and inflammation have been shown to change VWF binding altering the permeability of the blood brain barrier which can continue the cycle of inflammation and further damage brain tissues and pathways (Noubade et al., 2008; Ryu & McLarnon, 2009; Suidan et al., 2013; Zhang et al., 2002). Furthermore, differential editing in the following biomarkers can alter splicing profiles (NOVA1) (Irimia et al., 2012), the ability to repair DNA after oxidative stress (NEIL1) (Yeo et al., 2010), and cell differentiation and neurotransmitter release controlling synaptic plasticity (PCP4) (Wei et al., 2011). Differential editing plays a role in inhibitory signaling by editing GABRA3 at high levels in the adult brain. We found decreased editing in the PFC in both males and females, which can result in slower response to the GABA inhibitory neurotransmitter, thereby causing the cells to become more easily excitable (Rula et al., 2008). Excitatory signaling through kainate glutamate receptor GRIK2 can affect ion channel mediated signaling by differential editing seen in bipolar disorders and has been found dysregulated in neuro psychiatric disorders (Silberberg et al., 2012). Although we found significant difference in editing landscapes, future functional studies should explore, analyze and test the involvement of individual editing sites associated with symptoms of depression and suicide.

In the future, single cell-based transcriptome analysis can be used to identify nuances of differential ADAR editing landscapes, including between neural cell types within smaller brain regions and individual neural circuits. However, currently the accuracy of computational approaches of variant calling relies on a read depth greater than 15 (Ouyang et al., 2018) and most single cell datasets are currently performed using 10X genomics which is mostly below the threshold for accuracy. Since the majority of single cell RNA-seq experiments usually have read depths of no more than 10, they may not be as accurate in determining ADAR editing events. Furthermore, our approach to elucidating nuanced differential editing patterns and identifying relevant biomarkers can be extended to other conditions beyond neuropsychiatric disorders, particularly in the context of immune response, although further research is needed to directly connect clinical symptoms and observed phenotypic differences to the subtle changes in spatiotemporal editing landscapes.

We would like to note that although some of the samples had lower read depth, thereby potentially limiting our power to identify editing events, we were still able to identify differential patterns while controlling for false positives. We accomplished this by applying quality control measures for Guttman scaling, such as only including sites that were present in at least two samples within a condition and with a stack depth of 10 or larger. While we may have omitted some events with low editing frequencies, the reduction in the number of potential false positive events outweighs the impact of missing the few low-frequency editing sites. Our proposed statistic modeling approach that combines Guttman scaling with random forest modeling enables us to glean valuable insights even from relatively small samples sizes such as was used here however, we would like to advocate for the need of further studies with larger sample sizes to model more precise prediction of suicide. Moreover, since these editing profiles show differences between brain regions, there underscores the need for a less invasive way of measuring editing profiles such as through blood or saliva tests. In the future, to find clinically relevant biomarkers for diagnosis, prognosis or even therapy, editing profiles would need to be confirmed in blood and other bodily fluids.

## Conclusions

Our novel statistical model of quantifying Guttman scaling rank order and maximum likelihood scores with random forest modeling provides a way to take advantage of high-depth sequencing datasets, whether already available or newly obtained, to identify clinically relevant biomarkers. These biomarkers can provide insight into molecular mechanisms in differential brain regions underlying MDD and suicide. Additionally, these diagnostic, prognostic and therapeutic biomarkers once confirmed can be used in conjunction with current psychological assessments to determine risk for MDD and suicidal behavior. Here we leverage two previously published RNA-seq data sets to delineate ADAR editing landscapes and use these differential editing profiles for prediction of potential biomarkers for suicide and depression.

## Authors’ contributions

NMP conceived the project, conducted data analysis, and wrote the manuscript. RM contributed to conceptualization of analysis steps and data interpretation. HP conceived and supervised the project, assisted with data analysis, and wrote the manuscript. All authors read and approved the final manuscript.

## Acknowledgements

Not applicable.

