## Supplementary figures for "Differential ADAR editing landscapes in major depressive disorder and suicide"

**Supplementary Tables are available as separate XLSX file.**

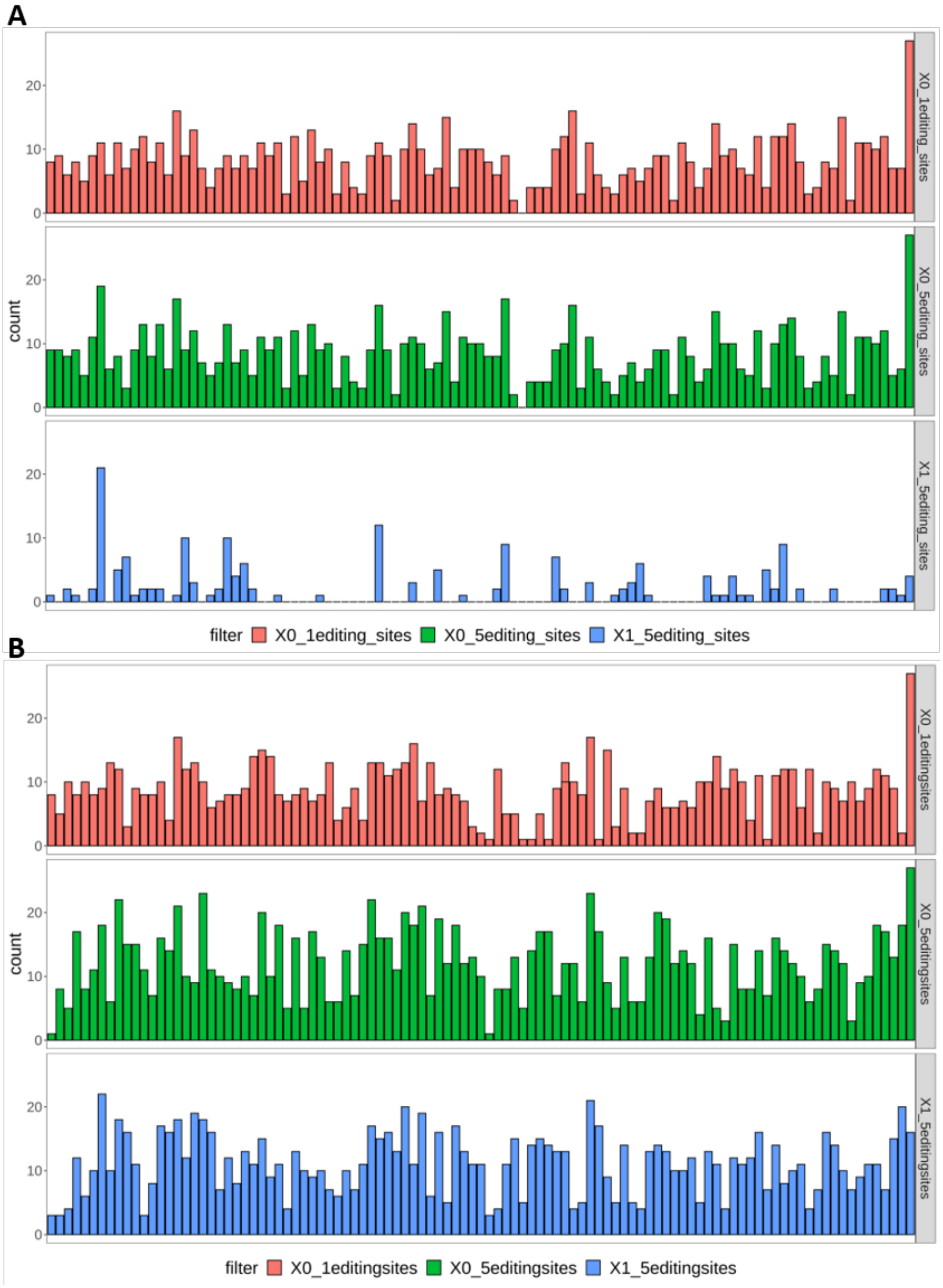
A
X0_1editing_sites
X0_5editing_sites
X1_5editing_sites
count
0
10
20
filter X0_1editing_sites X0_5editing_sites X1_5editing_sites
B
X0_1editingsites
X0_5editingsites
X1_5editingsites
count
0
10
20
filter X0_1editingsites X0_5editingsites X1_5editingsites

Supplemental Figure 1: Comparison of the frequency of A to G percent cutoff measures for using excitome sites for further analysis using Guttman Scaling. The excitome sites are along the X-axis in alphabetical order. COD is shown in A with red bars representing the changes in Guttman Scale rank order between no cut-off and 1% frequency cut off. Green is no cut-off compared to 5% frequency cut-off. Blue is 1% cut-off compared to 5% frequency cut-off. B shows the same comparisons but for MDD diagnosis. This shows there is over all a change in rank by 10 in both 1% cut-off and 5% cut-offs compared to no cut-off but it is uniform across sites.

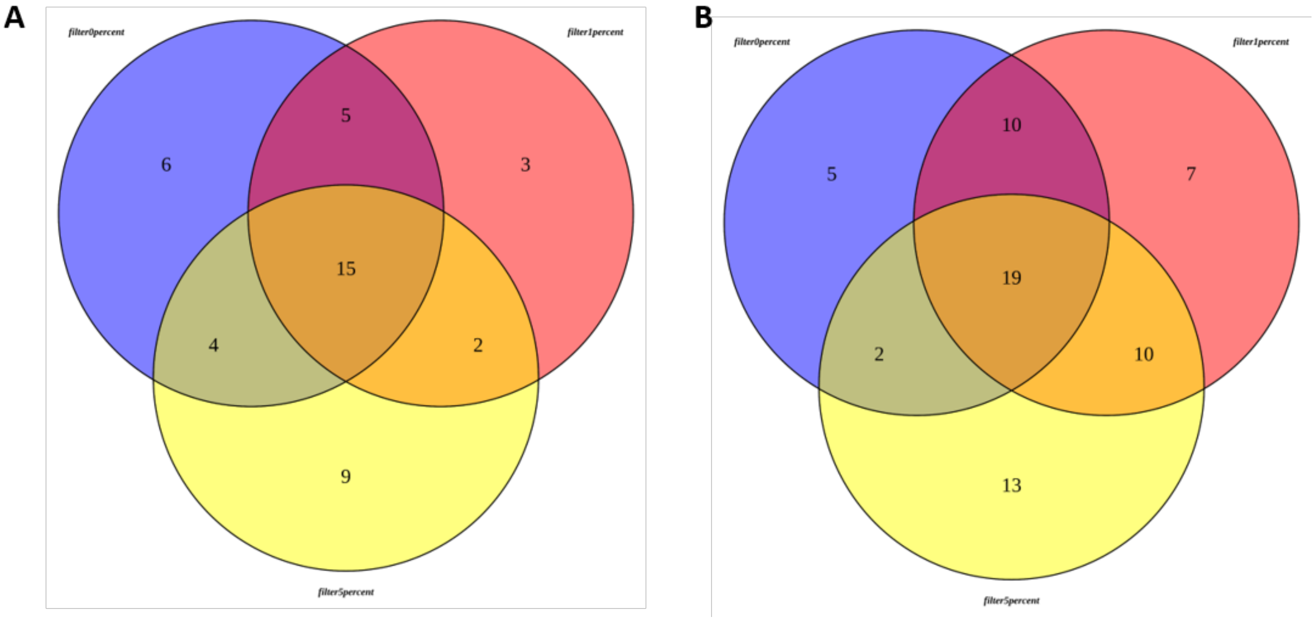


Supplemental Figure 2: Venn diagram illustrating that differences between different filtering cut-off showing that most variables used in our analyses remained the same regardless of the cut-offs, and that these variables included all the editing edits predicted to be potential biomarkers.

Supplemental Figure 3: Flow chart for fine-tuning random forest parameters

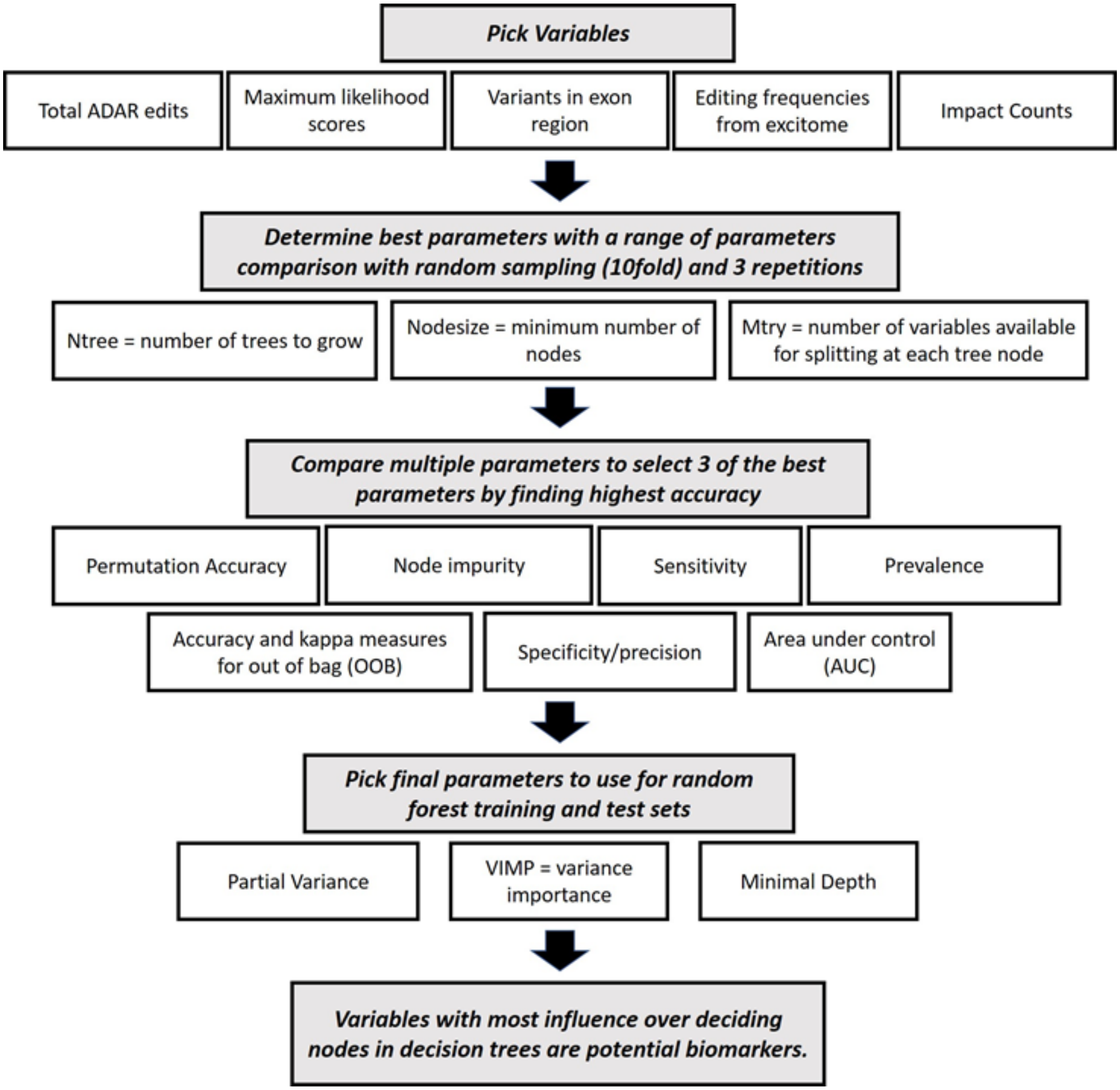


Supplemental Figure 3: Flow chart for complete classification random forest analysis including fine-tuning parameters, choosing between the best fine-tuned parameters after random sampling repetition based on specific criteria, and using multiple mathematical calculations to determine the top potential biomarkers.

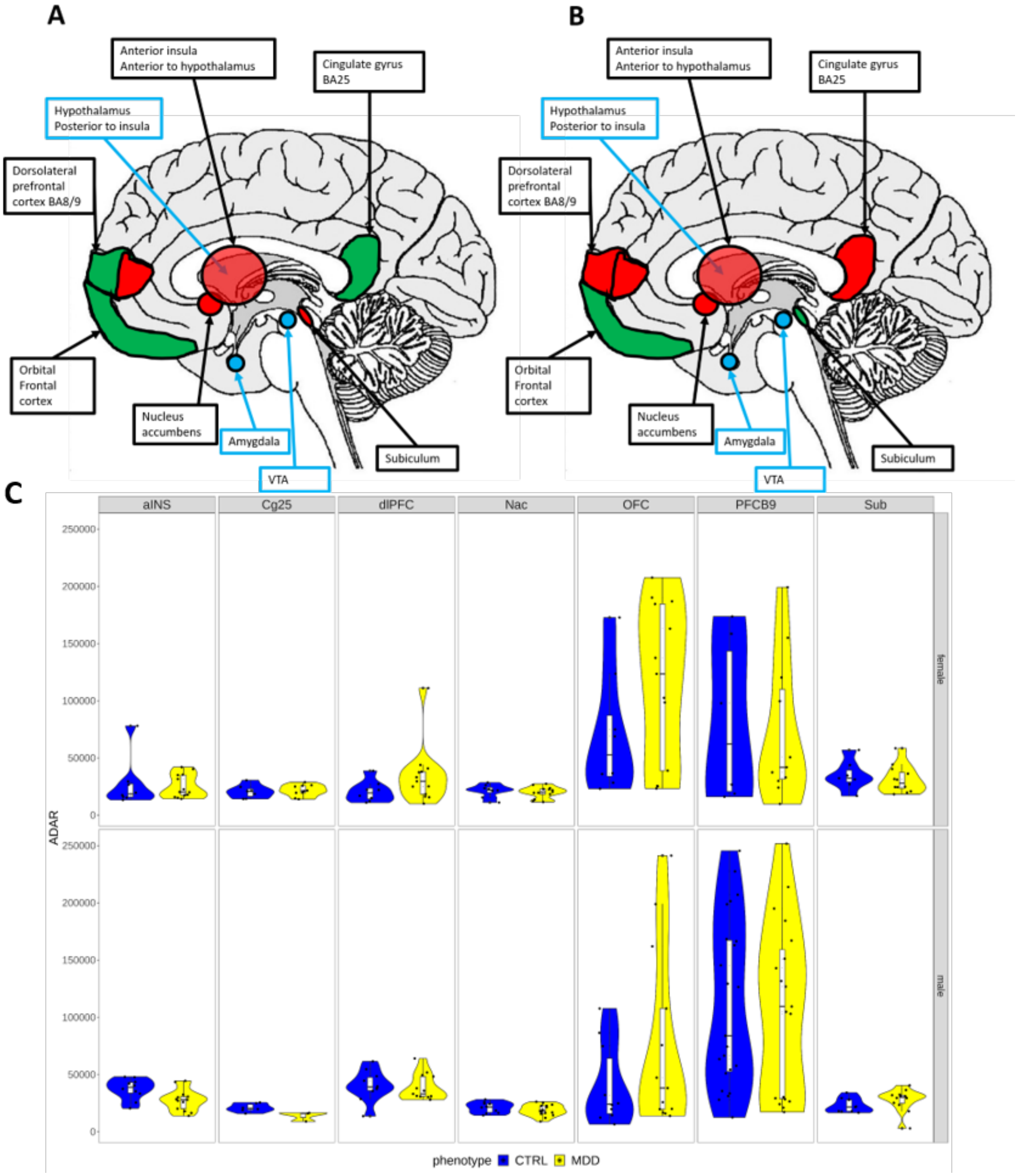
A
B
Anterior insula
Anterior to hypothalamus
Cingulate gyrus
BA25
Hypothalamus
Posterior to insula
Dorsolateral
prefrontal
cortex BA8/9
Orbital
Frontal
cortex
Nucleus
accumbens
Amygdala
Subiculum
VTA
C
aINS
Cg25
dlPFC
Nac
OFC
PFCB9
Sub
female
male
ADAR
250000
200000
150000
100000
50000
0
phenotype
CTRL
MDD

Supplemental Figure 4 Patterns of global ADAR editing changes across MDD diagnosis in females (A) and males (B). Increases in editing are shown in green, decreases in editing are shown in red, and no change is shown in yellow. Reference anatomy features shown in blue helps in the location of brain regions and are involved in pathways within studied regions. (C) Global differential ADAR editing patterns in MDD vary between brain regions in females and males, as well as between MDD and control (shown in yellow and blue in violin plots, respectively).

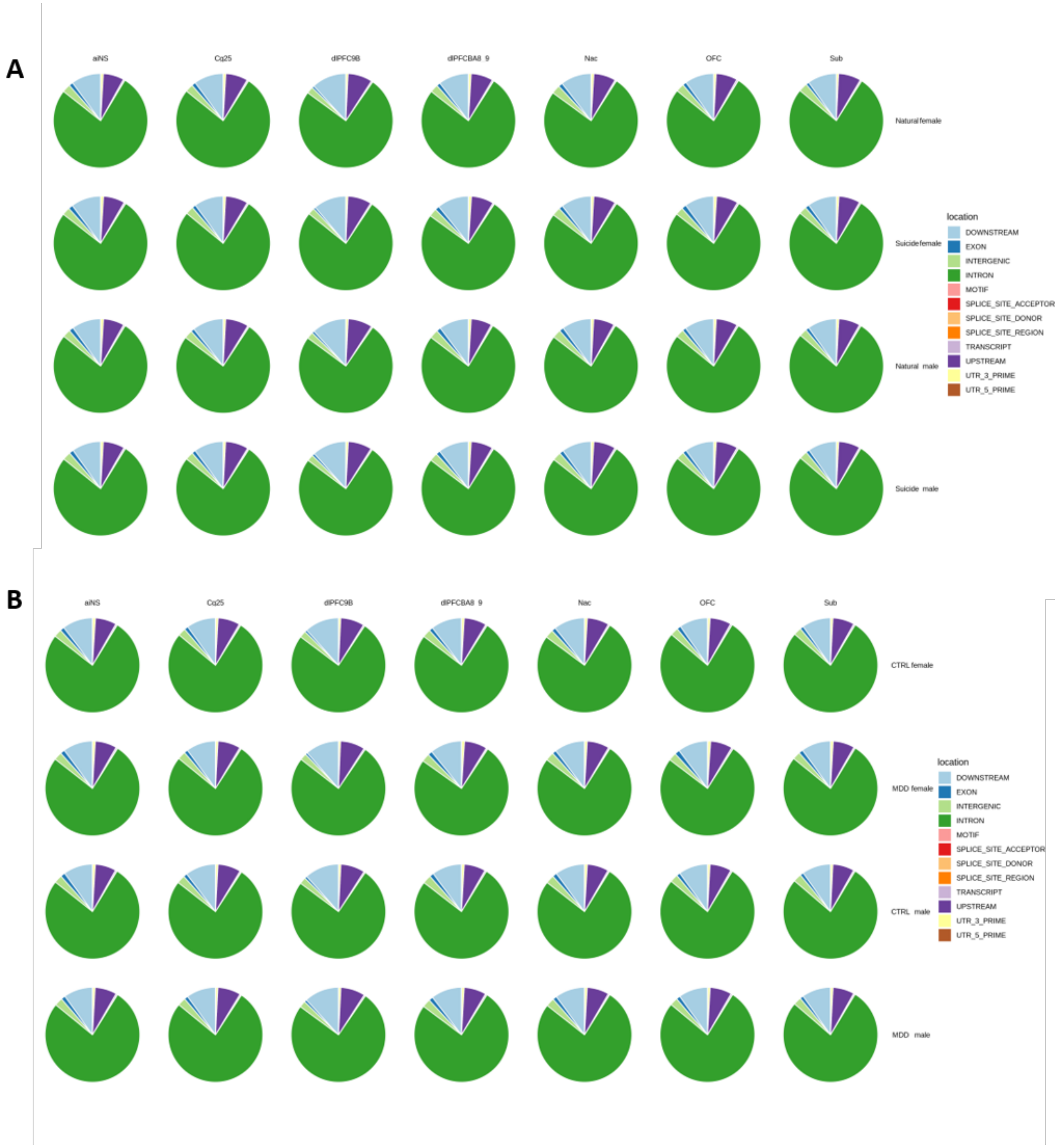


Supplemental Figure 5:
Location of global editing sites in COD (A) and MDD diagnosis(B). There is no difference between COD, MDD diagnosis, gender and tissue with most editing sites found in the intronic regions.

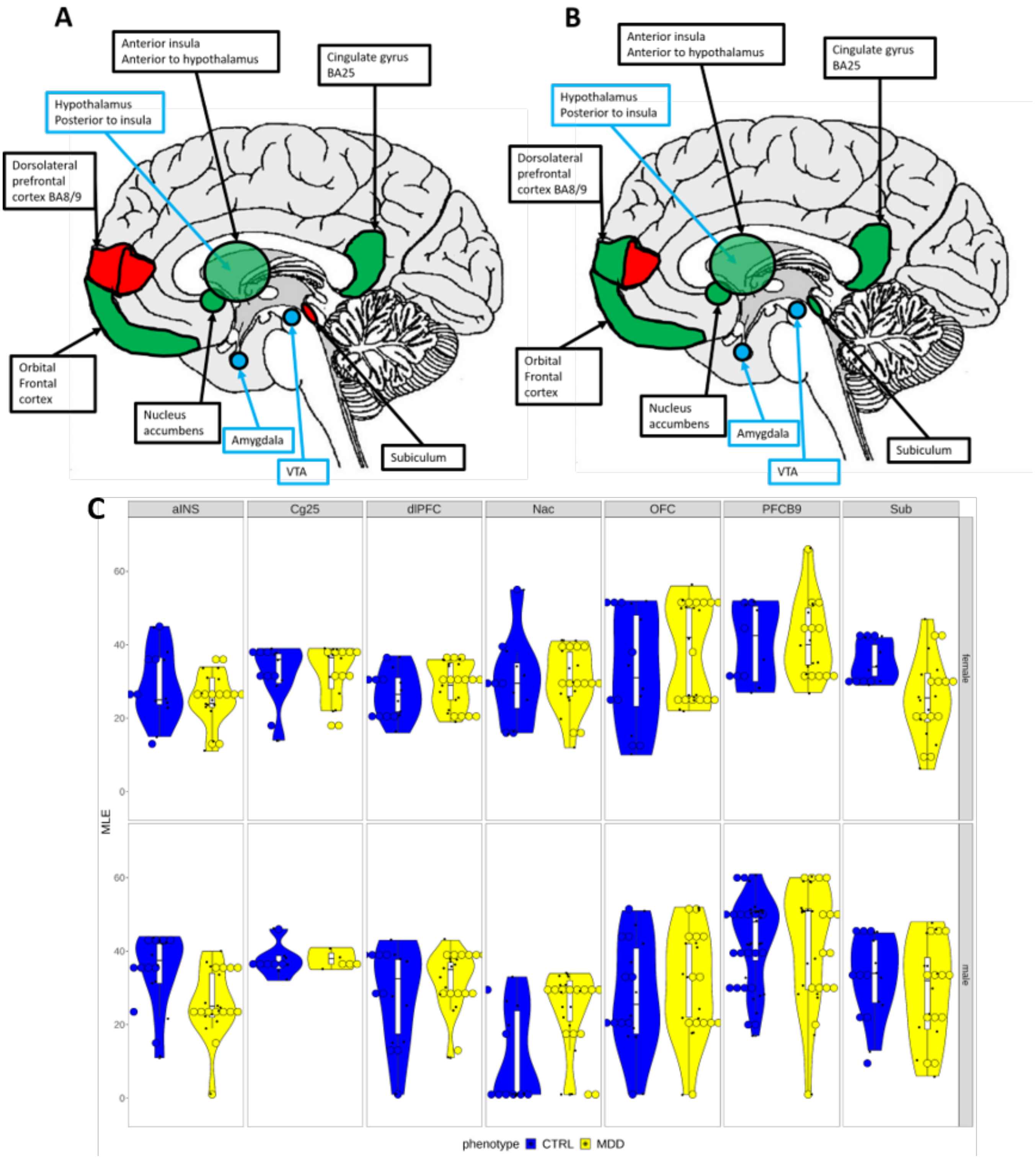

A
B
C
Anterior insula
Anterior to hypothalamus
Cingulate gyrus
BA25
Hypothalamus
Posterior to insula
Dorsolateral
prefrontal
cortex BA8/9
Orbital
Frontal
cortex
Nucleus
accumbens
Amygdala
VTA
Subiculum
aINS
Cg25
dlPFC
Nac
OFC
PFCB9
Sub
MLE
female
male
phenotype
CTRL
MDD

Supplemental Figure 6: Differences in Guttman maximum likelihood scores across MDD diagnosis in females (A) and males (B). Differential ADAR editing sites found in Guttman scale MLE scores in MDD are highlighted, with increases in editing shown in green, decreases in editing shown in red, and no change shown in yellow. Reference anatomy features shown in blue helps in the location of brain regions and are involved in pathways within studied regions. Similarly to the global differential ADAR editing patterns, Guttman scale MLE scores vary between brain regions in females and males in MDD, as well as between MDD and control (shown in yellow and blue in violin plots, respectively).

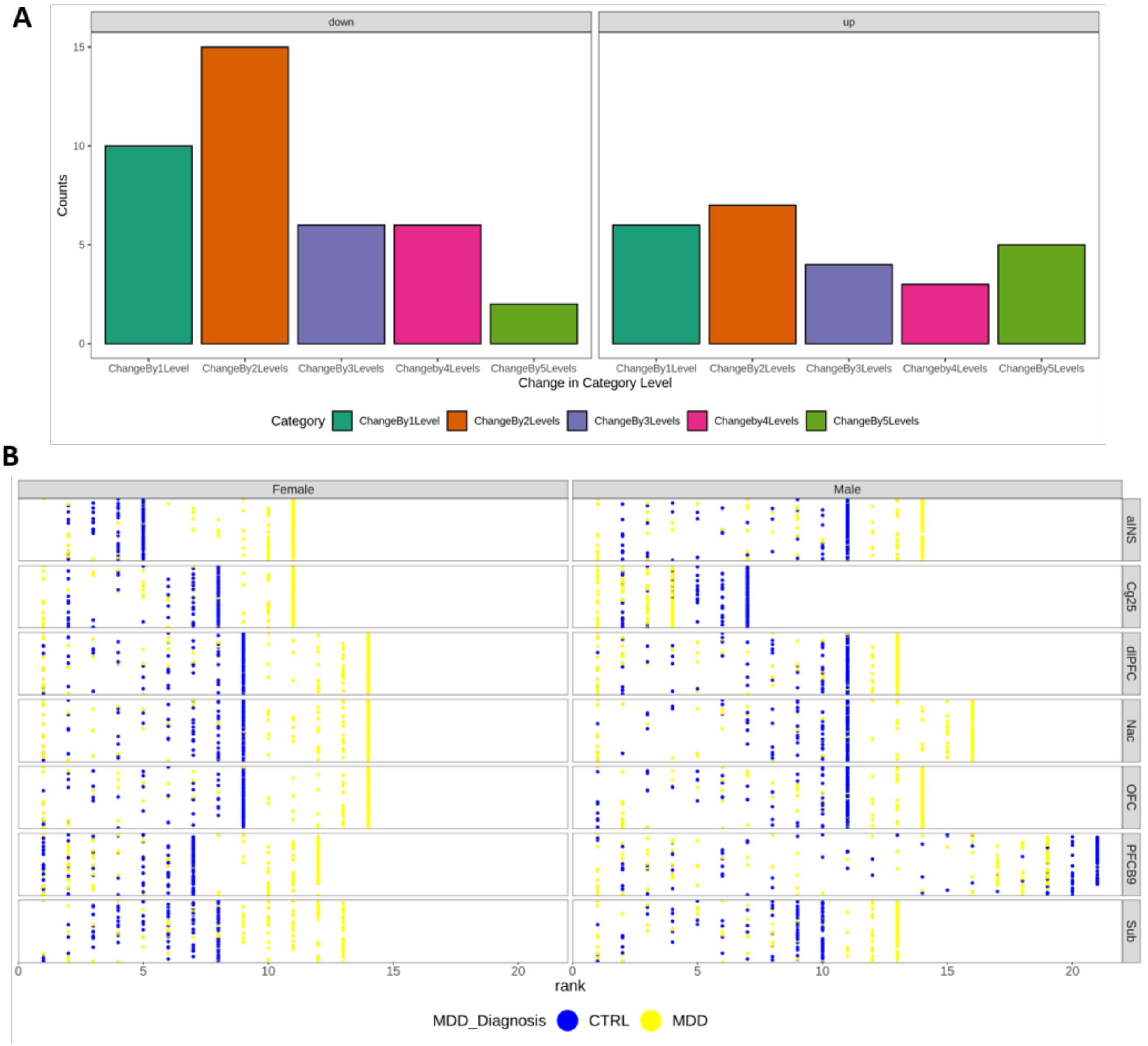


Supplemental Figure 7: Changes in Guttman scaling rank order determined by editing status of each site in MDD and non-MDD controls. Number of genes that change in Guttman rank order, whether up or down, from one rank change to five rank changes (A). Pattern of rank order as assigned by Guttman Scaling with non-MDD diagnosis shown in blue has slightly less variation in editing compared to MDD diagnosis shown in yellow. Females have more variation in editing sites indicating more editing with MDD diagnosis then males.

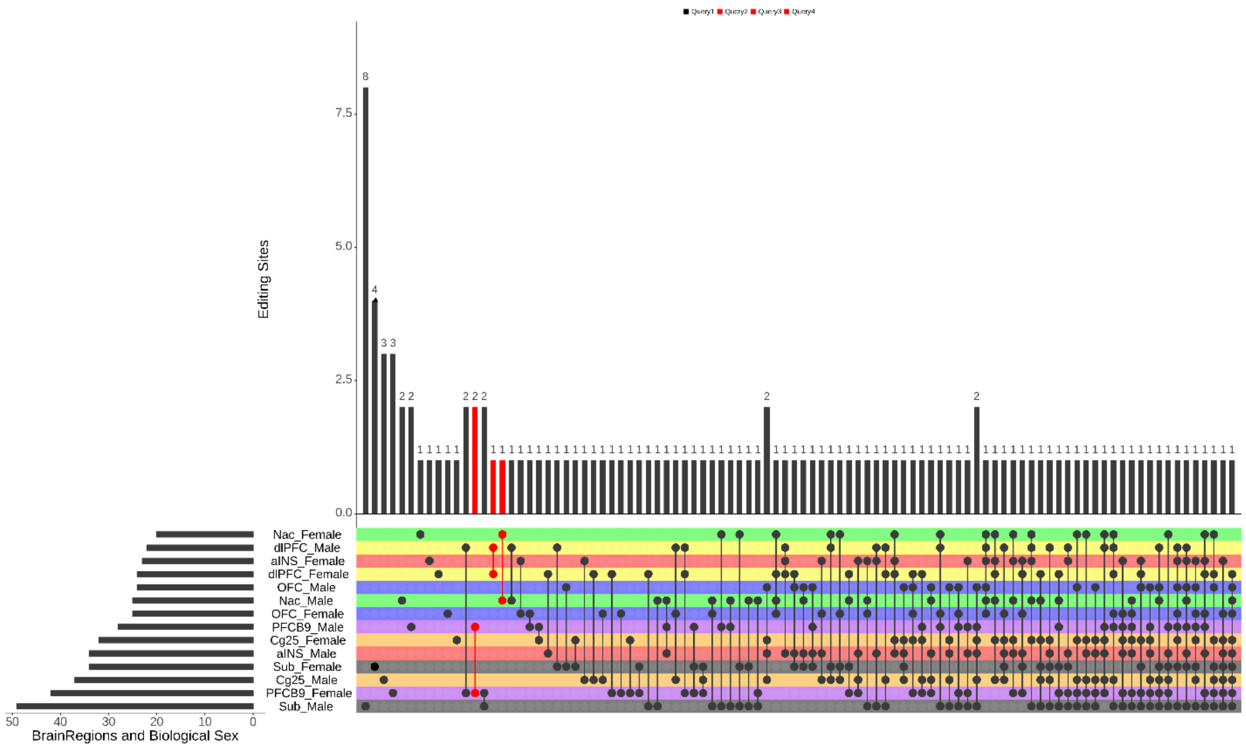


Supplemental Figure 8: Upset plot of genes within regions and sexes that shared similar changes in Guttman rank category editing status in MDD diagnosis.  The upset plot (Lex et al., 2014) shows each region in a different color, with each sex showing matching colors. The bars to the right show how many editing sites have changes to category, in order from the least amount of editing changes on the top with the most on the bottom. The bars across the top show how many editing sites are in each intersection. The dots in the colored bars represent intersections of the same editing sites in the region the dots are connected too. The connected dots shown in red represent the same region where both sexes share the same editing sites. For example, the first dots in red represent 2 editing in common between females and males in OFC which has 28 changes to editing sites in males and 41 in females. The second red dots represent 1 editing site in common between females and males in the dlPFC which has similar total number of changes just under 30. The last set of red dots shows 1 editing site in common between biological sexes in the Nac with just under 30 total editing sites in both biological sexes. Larger number of dots under the bar indicate higher number of regions that share changes in that editing site.

Supplemental Figure 9A:

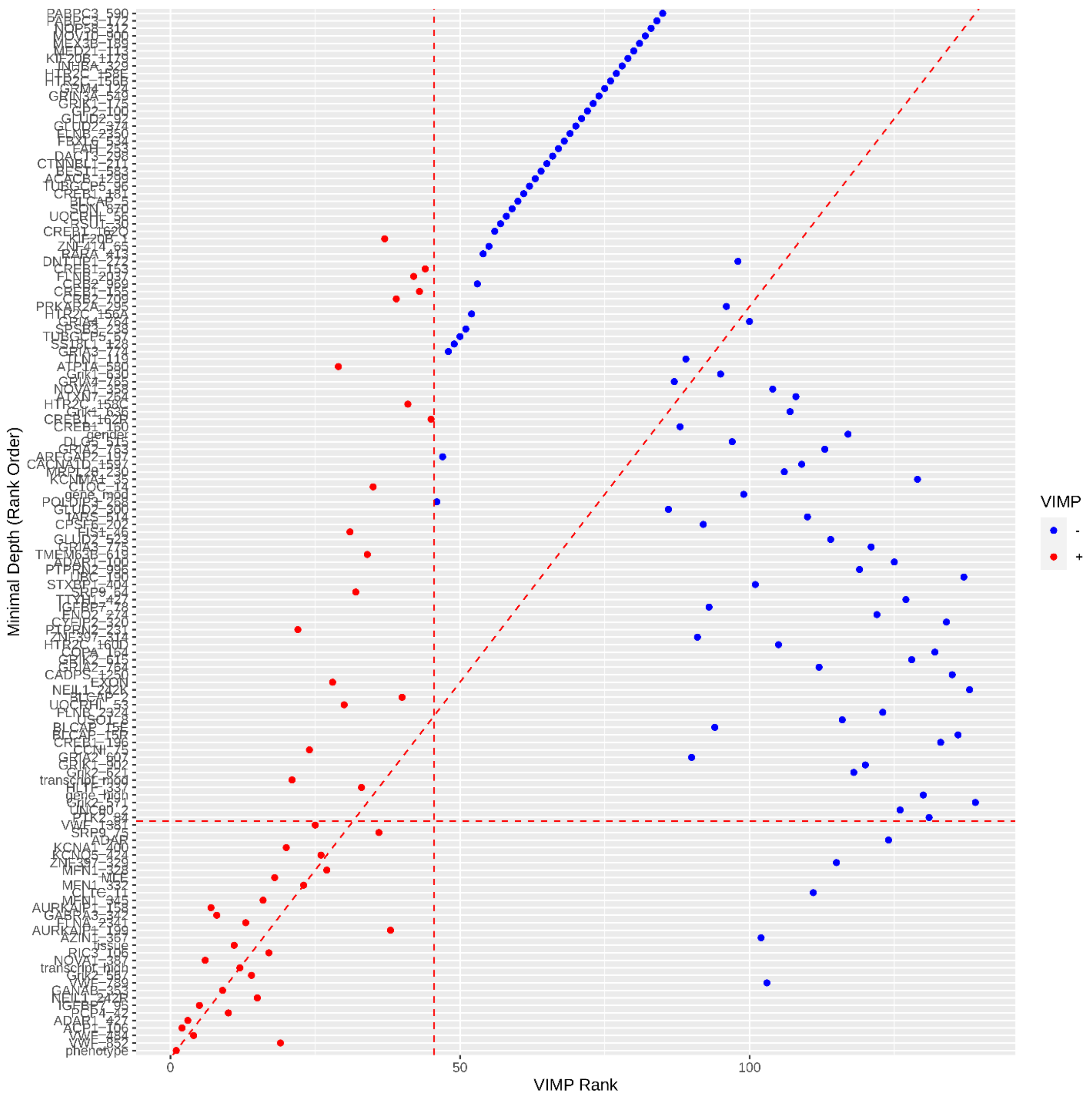

Supplemental Figure 9B:

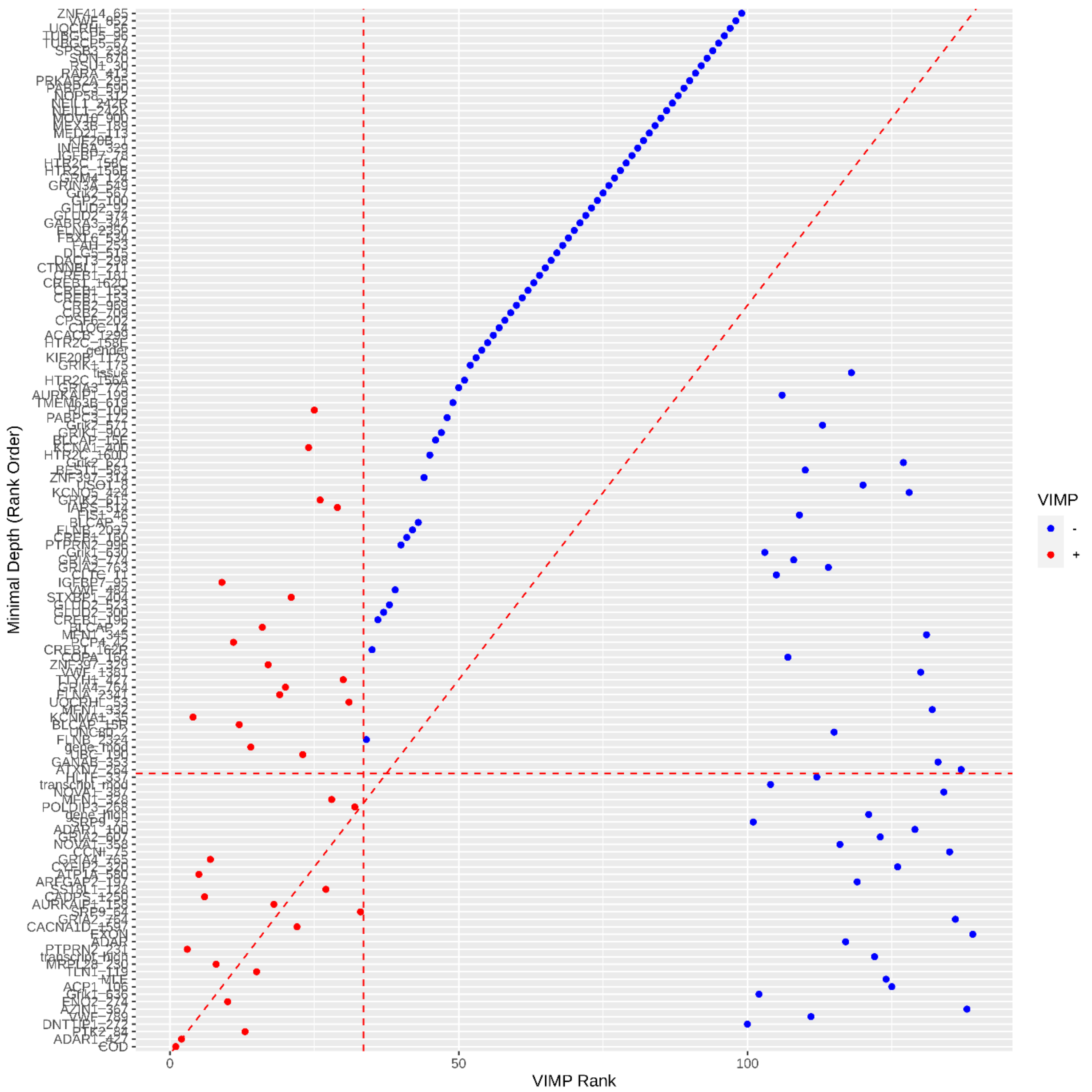


Supplemental Figure 9:Random forest top variants as determined by Variance importance (VIMP rank) and by minimal depth rank order (mindepth) show top candidates for potential biomarkers including all the variants found in partial variance analysis, but there were a few others that were near the high end of the cut-off for either VIMP or mindepth that were not found with variance importance and  thus were filtered out of the final biomarker list. This is true for both COD (A) and MDD diagnosis (B).

Supplemental Figure 10A

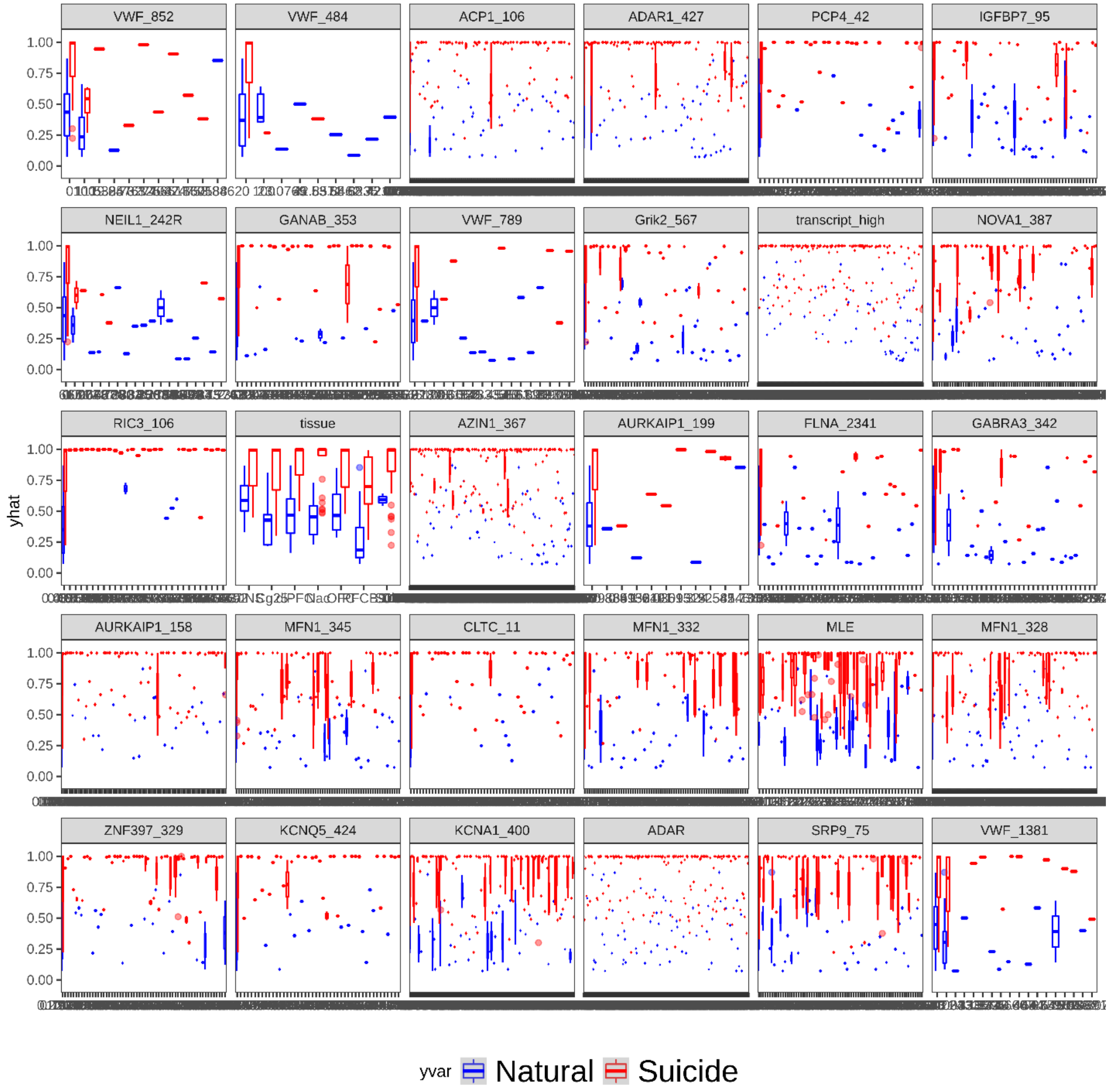

Supplemental Figure 10B

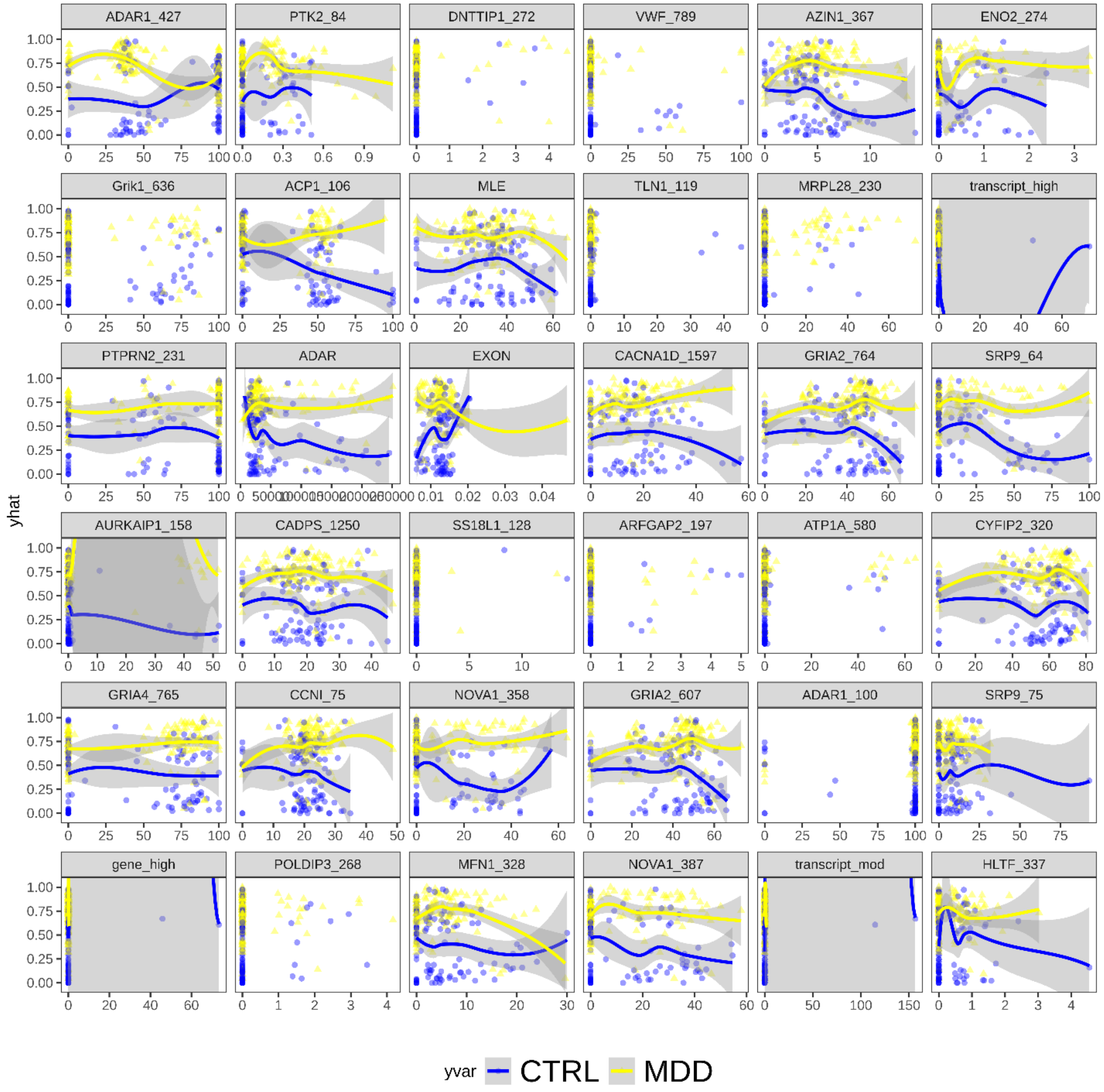


Supplemental Figure 10 Random forest top variants as determined by partial variance which were also found in VIMP/minimal depth comparison. Suicide random forest models (A) with control (natural COD) are shown in blue and suicide are shown in red, respectively. These individual panels show editing sites that are the most important in creating the random forest classification model. They show the variance between sites to be substantially different between COD, represented by the clustering of suicide cases with other suicides and controls with other controls (9A). This similar pattern is also seen in MDD diagnosis, with MDD diagnosis shown in yellow and control with no diagnosis shown in blue (9B). The top potential biomarkers are similar between COD and MDD diagnosis.

**Supplemental Methods**

In general, the larger the node size the fewer the variables used which results in smaller trees. Here we would assume a more complex, larger tree and start with smaller node sizes in comparison models. To ensure reproducibility and consistency between all random forest predictions, the seed was set to 7. When comparing mtry and nodesize in COD model, the highest accuracy was seen at nodesize 10 and mtry 140 (Supplemental Figure 11A). When comparing nodesize to ntree, we see the highest accuracy at nodesize 7 and ntree 50 (Supplemental Figure 11B). When ntree is set at 1500 and mtry is set at 140, the accuracy approaches 90% (Supplemental Figure 11C). The parameter training resulted in three potential parameter settings: (i.) nodesize 8, mtry 120 and ntree 1500, (ii.) nodesize 7, mtry 120, and ntree 50 and (iii.) nodesize 10, mtry 140, and ntree 1000. When comparing models of MDD diagnosis, the same parameter comparisons were made showing optimal settings of nodesize 6 and mtry 70 (Supplemental Figure 11D), nodesize 8 and ntree 300 (Supplemental Figure 11E), and ntree 50 and mtry 120 (Supplemental Figure 11F). The parameter training resulted in three potential parameter settings: (i.) nodesize 7, mtry 120 and ntree 50, (ii.) nodesize 8, mtry 50, and ntree 50 and (iii.) nodesize 5, mtry 70, and ntree 500.

To compare the three settings models to each other to find the best one, we used resampling without replacement (also in the caret R package) and the randomForest R package to run 10 potential random forest models and to calculate the accuracy of all the models. All missing data were replaced with the median of existing data. Importance of each variable was also calculated using two measures, (i). permutation accuracy (MeanDecreaseAccuracy), and (ii). Node impurity (MeanDecreaseGini), and accuracy and kappa measures using out of bag (OOB) error confusion matrix were used to determine the best parameter settings. The three model parameters for cause of death were highly similar, but the model with mtry of 140, nodesize of 10, and ntree of 1000 appeared to be slightly better. With MDD diagnosis, mtry 120, nodesize 7 and ntree 50 were the parameters with the highest accuracy as shown in Supplemental Table 9.

These models are parameters used for rfsrc R package to develop the model to predict biomarkers for suicide risk and those for MDD diagnosis, including: (i.) maximum likelihood scores (calculated with Guttman scale), (ii.) number of ADAR variants found in the exon regions (calculated by snpEff), (iii.) total number of ADAR variants (calculated with ExToolset) and (iv.) editing frequencies of each editing site in the excitome (calculated with ExToolset). The importance of each variable on the model was calculated using two parameters minimal depth set to a cut-off determined by mean of the minimal depth distribution and VIMP in rank order set to a cutoff of 40. The variables that meet the cut-offs for both will be the final biomarkers. Accuracy, misclassification rate, true positive rate, false positive rate, true negative rate, precision, prevalence and null error rate were calculated from each cause of death and MDD diagnosis models.

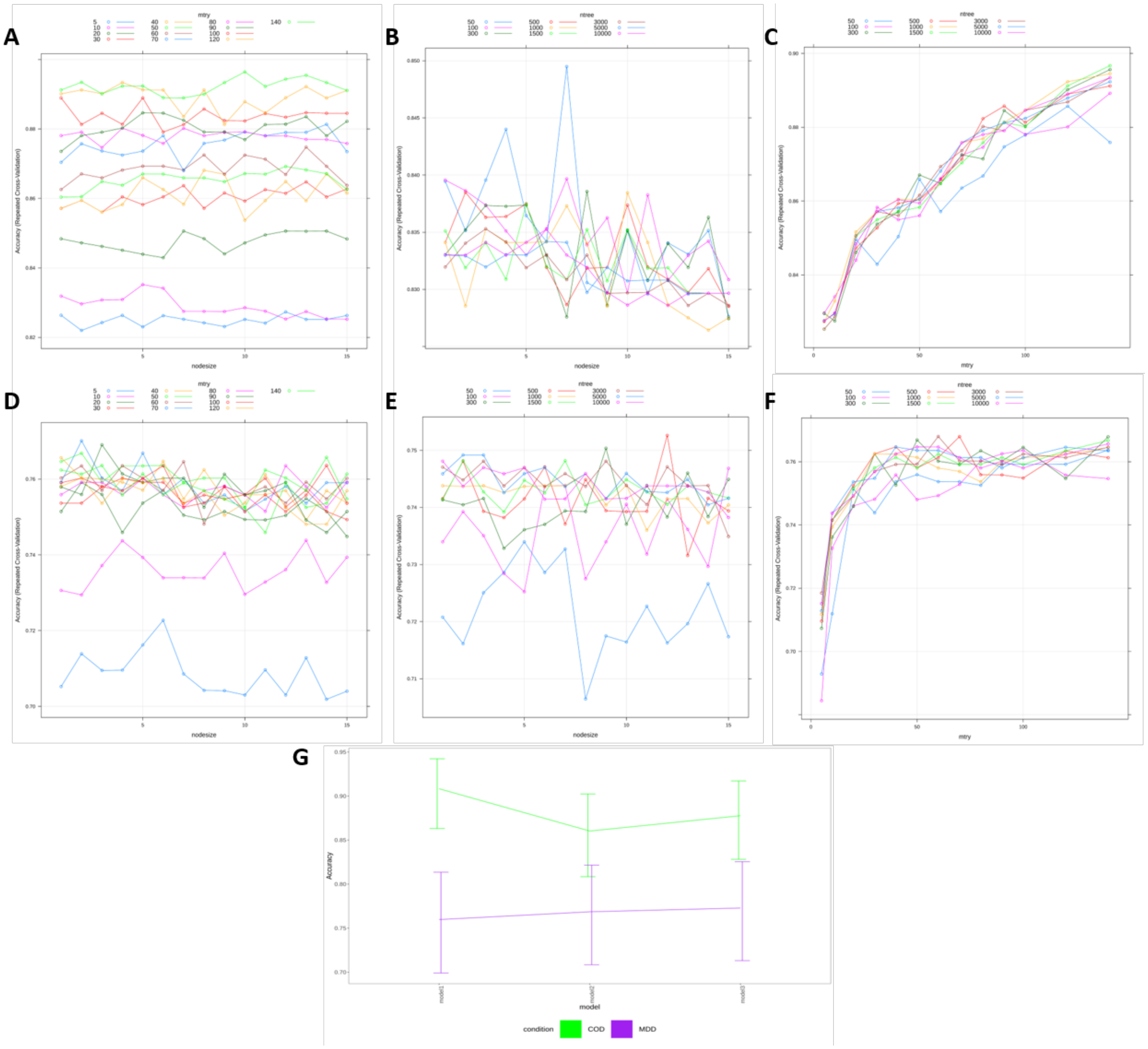

Supplemental Figure 11: Random Forest Parameter tuning parameters, showing varying model parameters used to fine-tune random forest classification. By finding the highest accuracy rate for each parameter used to construct trees including mtry (A), node size (B) and ntree (C) for COD and the same for MDD (D-F). With these 3 parameter models created for each COD (green) and MDD (purple), we can then use the accuracy found in random resampling techniques to find the best set of model parameters to run the random forest classification modeling with.
